# Optic flow helps explain gulls’ altitude control over seas

**DOI:** 10.1101/569194

**Authors:** Julien R. Serres, Thomas J. Evans, Susanne Åkesson, Olivier Duriez, Judy Shamoun-Baranes, Franck Ruffier, Anders Hedenström

## Abstract

For studies of how flying animals control their flight, seabirds are of particular interest to track with a biologger because they forage offshore where the visual environment can be simply modeled by a flat world textured by waves. This study suggests that optic flow can explain gull’s altitude control over seas. In particular, a new flight model that includes both energy and optical invariants (called the *ventral optic flow regulation*) explain the dynamics of gulls’ altitude control during offshore takeoff and cruising flight. A linear statistical model applied to 352 flights from 16 individual lesser black backed gulls (*Larus fuscus*) gave a strong correlation between wind assistance and gulls’ altitude. Thereafter, an optic flow-based flight model was applied to 18 offshore takeoff flights from 1. individual gulls. By introducing an upper limit in climb rate in a non-linear first order parametric model on the gull’s elevation dynamics, coupled with an optic-flow set-point, the predicted altitude gives an optimized fit factor value of 63% on average (min value: 30%, max value: 83%) with respect to GPS data. We conclude that the optic-flow regulation principle (here running close to 25°/*s*) allows gulls to adjust their altitude over sea without having to directly measure their current altitude.

## 1. Introduction

Understanding how a bird decides to fly at a given altitude during a specific manoeuver is a difficult task because it is strongly dependent on the atmospheric conditions and flight capacity of the bird (see review [59]). Seabirds such as albatrosses and petrels flying close to the sea surface take advantage of the logarithmic increase in wind speeds to support dynamic soaring [50,52,53, 66], which works only at very low altitudes from ca. 0-10 m (see e.g. Fig. 5 in [56]). Birds flying by flapping flight at low altitudes over the sea could also use this windspeed gradient to reduce their transport costs. Under tailwinds, birds should fly higher where wind speed is high, while under headwinds birds should fly lower where wind speed is low. In terms of energy, a bird minimizing its transport cost should adjust its airspeed with respect to wind by increasing it in headwinds and decreasing it in tailwinds [26, 48]. This prediction comes from a U-shaped function between power required to fly and airspeed, which defines characteristic speeds for achieving minimum power *V_mp_* and maximum range *V_mr_*. During migratory [38] and homing flights [33] birds utilize wind assistance to minimize the transport cost and adjust airspeed accordingly to fly at the wind dependent *V_mr_*.

Groundspeed is the combined effect of airspeed and wind speed (actually the airspeed and wind vectors). Wind assistance alone cannot be used by the bird to select a given groundspeed and a flight altitude. The altitude could be set by surrounding visual information seen by the bird. A bird can access information about its own motion with respect to its surrounding environment via the optic flow field through its early visual processing [4], as flying insects do in similar situations [4, 58]. The optic flow field perceived by an agent (a flying insect, a bird, or a human) is particularly dependent on the structure of the environment [19, 35, 45, 67]. Optic flow can be defined by a vector field of the apparent angular velocities of objects, surfaces, and edges in a visual scene caused by the relative motion between the agent and the scene (Fig. 1). The translational optic flow component is particularly interesting for birds positioning in space because it depends on (i) the ratio between the relative linear groundspeed of an object in the scene with respect to the bird, and (ii) the distance from obstacles in the surrounding environment. Consequently, optic flow requires neither groundspeed nor distance measurement, which is particularly useful to explain how birds perceive the world because birds are likely unable to sense directly their own groundspeed nor the 3D structure of the environment in which the binocular vision plays a minor role [39].

**Figure 1.**
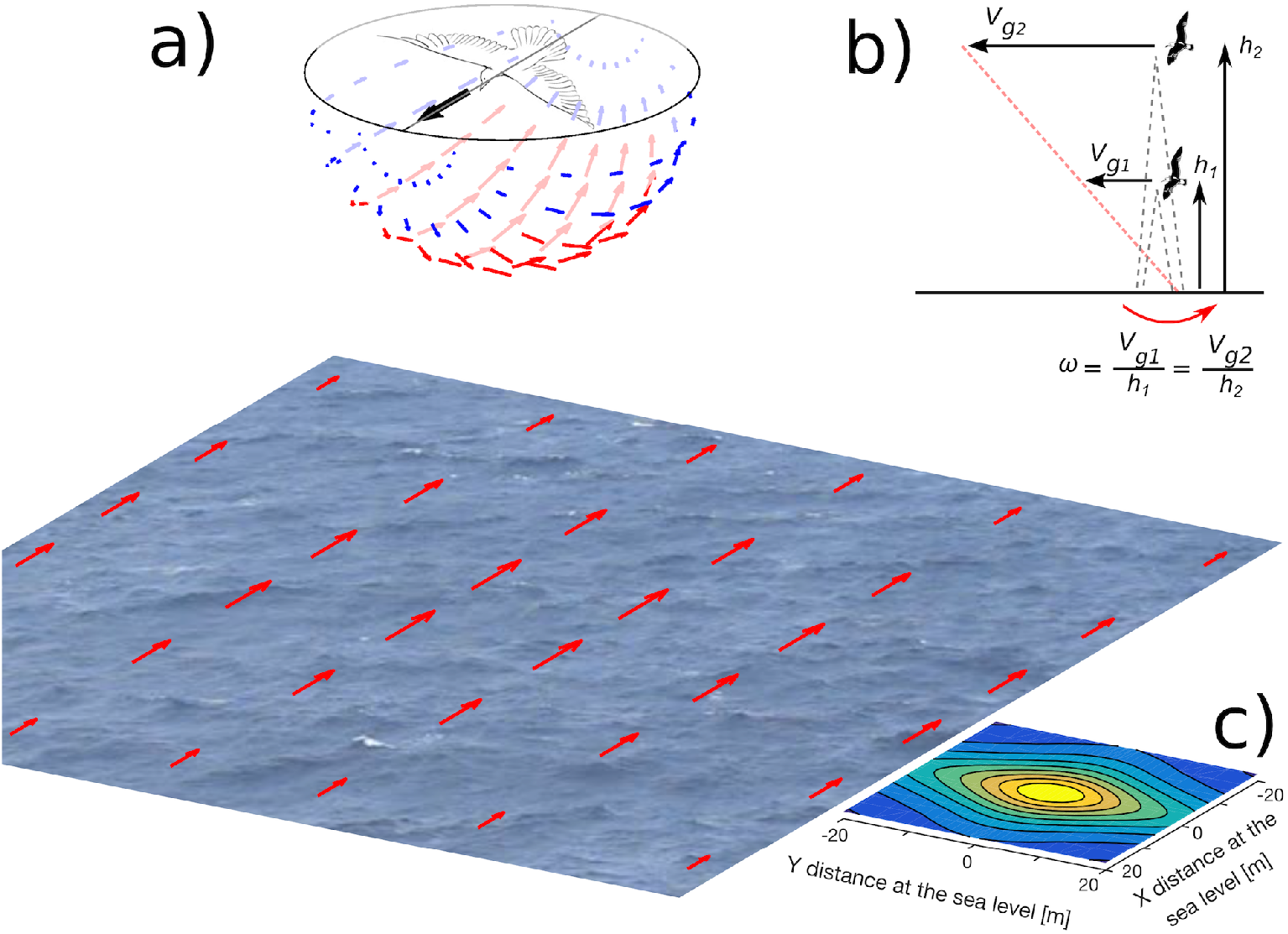
(a) A gull flying over the sea generates a vector field of optic flow. Such a vector field is perceived by a gull based on the contrasts created by waves and white-crested waves (also called white-horses). Inspired by [19]. (b) The magnitude of the vector of the optic flow, *ω*, is determined by the gull’s groundspeed, *V_g_*, and its altitude, *h*. If *ω* is held constant by adjusting the altitude, *h* will always tend (through the bird dynamics) to be proportional to *V_g_* (only a linear combination -red dashed line-between h and *V_g_* is asymptotically possible). (c) Optic flow magnitude in the ventral field of view at 10m-height where the magnitude of the ventral optic flow 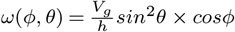 is projected at the sea level with *ϕ* the azimuthal angle and *θ* the elevation angle. (The magnitude of vertical optic flow is the maximum and is 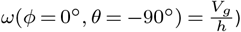

During flight manoeuvers, various optic flow parameters (such as the magnitude, the direction, the focus of expansion, the time-to-contact of optic flow) can be collected by birds to control their lateral position in straight tunnels (in budgerigars [7]), to decrease their speed in a converging tunnel (in budgerigars [57]), to plunge into water (in gannets [36]), to hover (in hummingbirds [20, 54]), and finally to land (in hawks [12] and in hummingbirds [37]).

In this study, we address the question of how seabirds control their altitude during offshore takeoffs and cruise flights with respect to wind. Here, two working hypotheses were compared about altitude control:

- a first hypothesis based on a direct measurement and regulation of optic flow that adjusts the altitude, and,
- a second hypothesis based on a direct measurement of the barometric pressure that directly regulates the altitude itself.

To test these alternative hypotheses, a statistical analysis of 352 flights comprising 16 individual lesser black-backed gulls (*Larus fuscus*) in various wind conditions were conducted.

Then, 18 offshore takeoffs followed by a cruise flight were analyzed by taking into account morphological parameters from 9 individual gulls.

## 2. Operating point in flight in terms of speed and altitude: a theoretical approach

### (a) How is bird speed deducted from aeraulic effects?

The relationship between:

– the bird’s ground speed *V_g_*
– the bird’s airspeed *V_air_*
– the wind speed *V_w_* is given by equation (2.1):

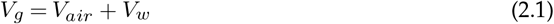

The basis for deriving predictions about bird flight is the so-called flight mechanical theory, which combines the relationship between power output *P* and airspeed *V_air_* in flapping flight as follows:

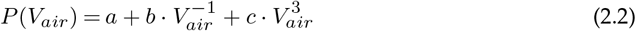

where *a, b*, and *c* represent various physical, morphological and physiological properties of the bird and air [47, 49, 51]. If the objective is to minimize the energy cost per unit distance (i.e., cost of transport), the optimal flight speed is the maximum range speed *V_mr_* [26, 47]. The maximum range speed *V_mr_* is obtained from the U-shaped power curve [24, 27, 51] by the condition:

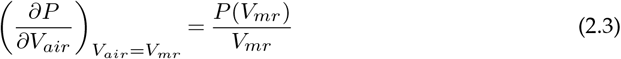

Indeed, a gull’s homing flight is similar to a migratory flight, in that it is assumed that the flight’s objective is principally for transportation, as opposed to outbound foraging flights when the bird is likely also searching for food. Seabirds’ homing flight over the sea is therefore a relatively straight path between two locations. During transport flight gulls are expected to minimise overall energy expenditure or time, thus cost of travel per unit distance should be minimised rather than instantaneous energy expenditure. If minimising the cost of travel per unit distance birds will travel at maximum range speed (*V_mr_*) not minimum power airspeed (*V_mp_*). *V_mr_* refers to *V_air_* rather than *V_g_*. If a bird experiences a tailwind, its cost of travel per unit distance decreases, thus *V_mr_* also declines. Conversely under headwinds *V_mr_* increases. In a recent work, it was analyzed how lesser black-backed gulls (and guillemots) modulate their airspeeds in relation to winds [17]. It was found that gulls increased airspeeds under headwinds and decreased airspeeds under tailwinds [17], and similar behaviour has been observed during longer distance homing flights [42]. These results suggest that gulls are flying at *V_mr_* rather than *V_mp_*, since *V_mp_* should not be affected by winds like *V_mr_* [17].

### (b) Optic flow vector field

Consider a bird flying over the sea, assumed as flat in the optic flow calculation, then based on groundspeed *V_g_* only (neglecting vertical speed *V_z_*) the magnitude of the ventral optic flow field *ω* can be expressed as follows:

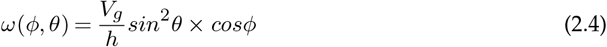

with *h* the altitude, *θ* the elevation angle and *ϕ* the azimuth angle.

The magnitude of the ventral optic flow field is plotted in Fig. 1a with the projection of its elevation and azimuth angles over the sea. The larger projection of vector magnitude of optic flow over the sea is shown using a contour plot in Fig. 1c in the case of a bird flying at a height of 10 m. The bird may be able to perceive the optic flow maximum from a non-negligible area of its field of view (Fig. 1c). The maximum magnitude of the ventral optic flow is always vertically downwards from the bird in the direction of the sea:

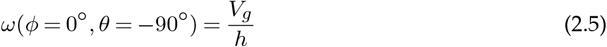

### (c) How the model predicts the bird’s flight height from the ventral optic flow regulation principle

The ventral optic flow regulation principle tends to keep constant the vertically downward optic flow whatever the speed or height of flight by adjusting the altitude [18, 55]. Here, it introduces this asymptotic proportionality relationship for birds: the bird’s height of flight *h* will always tend (through the bird dynamics) to be proportional to the bird’s ground speed *V_g_* (Fig. 1b) as:

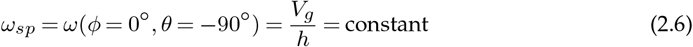

where *ω_sp_* is the ventral optic flow set-point. Besides, the wind profile power-law is often used to estimate the horizontal wind speed [31] as follows:

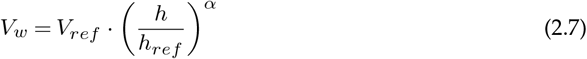

with the parameter *α* is the power-law exponent (that is usually specified as a function of stability as well as the roughness of the surface 0<*α*<1 (here over seas *α* = 0.11 see [30]), the speed *V_ref_* as being the wind speed at a reference height *h_ref_* (10m). By combining (2.6) and (2.7) into (2.1), we obtain:

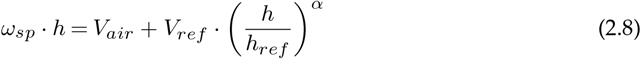

To find the bird’s steady-state flight height *h* reached during a takeoff as function of the wind profile, it requires to solve the equation *f*(*h*) = 0 with the function f defined as follows:

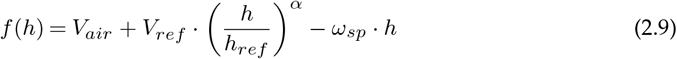

In the variation table of the function *f*(Tab. S1), we observe that only one unique altitude *h* exists, enabling *f*(*h*) = 0 during an offshore takeoff manoeuvre. We can therefore conclude that both the minimisation of the rate of energy consumption and regulating the ventral optic flow enable a bird to fix both its groundspeed and its altitude above the sea. The bird’s steady-state flight height *h* cannot be considered as a “target flight height” or a “desired flight height”, but as an “optimal flight height” because the bird’s altitude is adjusted as a function of the wind conditions (higher under tailwinds but lower under headwinds) and thereby maximizing positive effects as well as minimizing adverse effects of the wind gradient.

## 3. Gulls’ trajectory recording

16 lesser black-backed gulls (*Larus fuscus*) were GPS tracked from their breeding colony on Stora Karlsö island, Sweden (17.972° E, 57.285° N) during May to September of 2013-2015. The island is a small offshore island (2.5 km^2^) located in the western central Baltic Sea, sited 7 km west of the much larger island of Gotland (Fig. 2a). During breeding the gulls perform central-place foraging trips [46], flying out from their island to forage, either at sea or on land [32].

**Figure 2.**
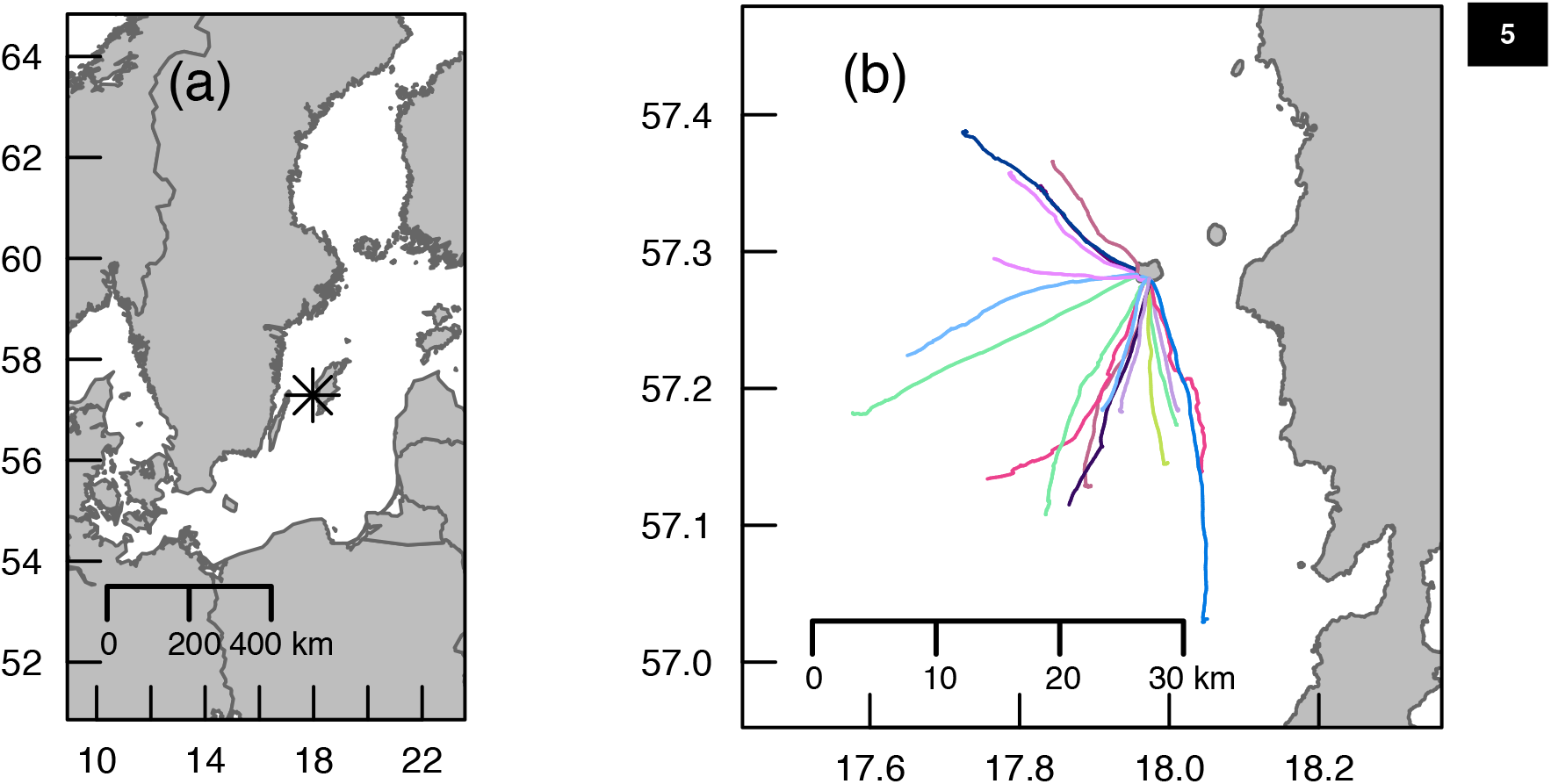
(a) Study location of the island of Stora Karlsö (indicated by *asterisk), Baltic Sea, Sweden. (b) From this site Lesser Black-backed Gull (*Larus fuscus*) inbound flights were tracked with GPS (18 flights from 9 individual gulls, coloured lines).

Gulls were caught during late incubation (late May) using walk-in traps set over their nests. They were weighed and sexed from morphological measurements [11] or genetically [23] from a few breast feathers taken at capture. An 18 g solar-powered UvA-BiTS GPS tracker with remote download capacity [9] was mounted using either a full body or wing harnesses [64] constructed of tubular Teflon™ ribbon (Bally Ribbon Mills 8476-.25”) (full tagging procedure given in [32], see Fig. 3). Data were downloaded and programs uploaded to the GPS devices remotely using a network of four antennas providing good coverage of the colony area. GPS tracking was continuous though the location intervals varied depending on the requirements of parallel studies (e.g. [32]). At a 6 seconds interval on a white stork (*Ciconia ciconia*) on its nest, it was quantified a mean altitude error of 2.77 m and a mean speed error of 0.02 m/*s* of the UvA-BiTS GPS tracker [9].

**Figure 3.**
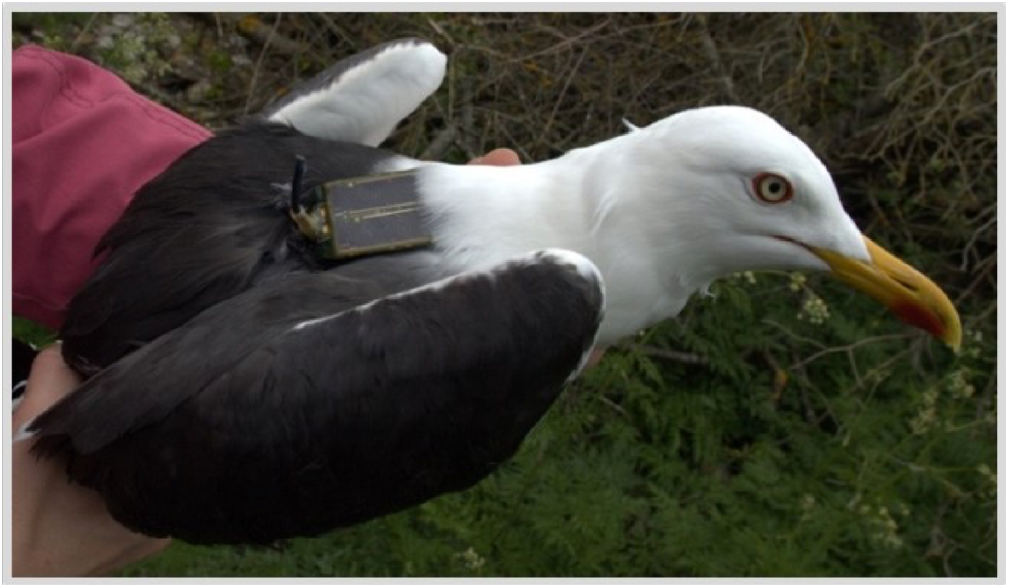
Lesser black-backed gull (*Larus fuscus*) equipped with an 18 g solar-powered UvA-BiTS GPS tracker (see [9] for GPS tracker details). Photographic credit: the authors.

The continuous GPS tracks were segmented into foraging trips and within these, sections of continuous flight, with the final flight of a foraging trip considered a homing flight, as the gulls returned from presumed foraging at sea (only marine trips were used in this study, c.f. [32]) to the island colony. 18 takeoffs from 9 individual gulls with high resolution data were selected (i.e. 10 or 15-second intervals), and we selected only takeoffs reaching a steady-state altitude - i.e. not those with a constantly fluctuating altitude. In addition, the final altitude had to be greater than 10 m with variation in altitude during the ascent until reaching a steady-state altitude. Flight GPS points were annotated with wind data extracted from a global weather model, ERA-interim data [13] provided by the European Centre for Midrange Weather Forecasts (ECMWF, http://www.ecmwf.int/en/research/climate-reanalysis/era-interim), which gives variables at 3-hour intervals and is gridded with a spatial resolution of approximately 79 km. These were extracted using the environmental-data automated track annotation (Env-DATA) system [14] hosted by MoveBank (http://www.movebank.org/).

## 4. Full flights’ dataset analysis: statistical model

The dataset here includes all inbound (returning to the island colony) over sea flights by the lesser black-backed gulls (383 flights, 16 gulls). The dataset is composed of median altitudes *h* calculated per flight, median wind speed measured at 10m-height (from ECMWF data), *V_ref_*, and the gull identifier. After excluding the flights endowed with a median altitude below zero meters, the data comprise 352 observations of 16 individual gulls.

A nonlinearity of wind profile power law (2.7) was introduced to estimate the wind speed *V_w_*(*h*) experienced by gulls at their median altitude *h* calculated per flight. A linear mixed effect model was designed using *lmer* in R software for the ordinates (*β_i_* is the constant random effect) as follows:

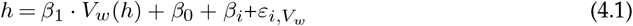

with the regression parameters: *β*_1_ = 2.2707 and *β*_0_ = 32.0016. The Kenward-Roger corrected F-test was used to calculate the significance level of the linear mixed model (ndf:1, ddf: 347.89, Fstat: 37.722, p.value: 2.2286·10^−9^, F.scaling: 1). The parameter *β*_1_ was highly significant (Fig. 4). Using the coefficient *β*_1_ = 2.2707, an identification of the ventral optic flow set-point *ω_sp-lmer_* can be performed using the equation (2.8) that includes the wind profile power law as follows:

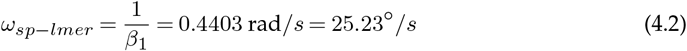

**Figure 4.**
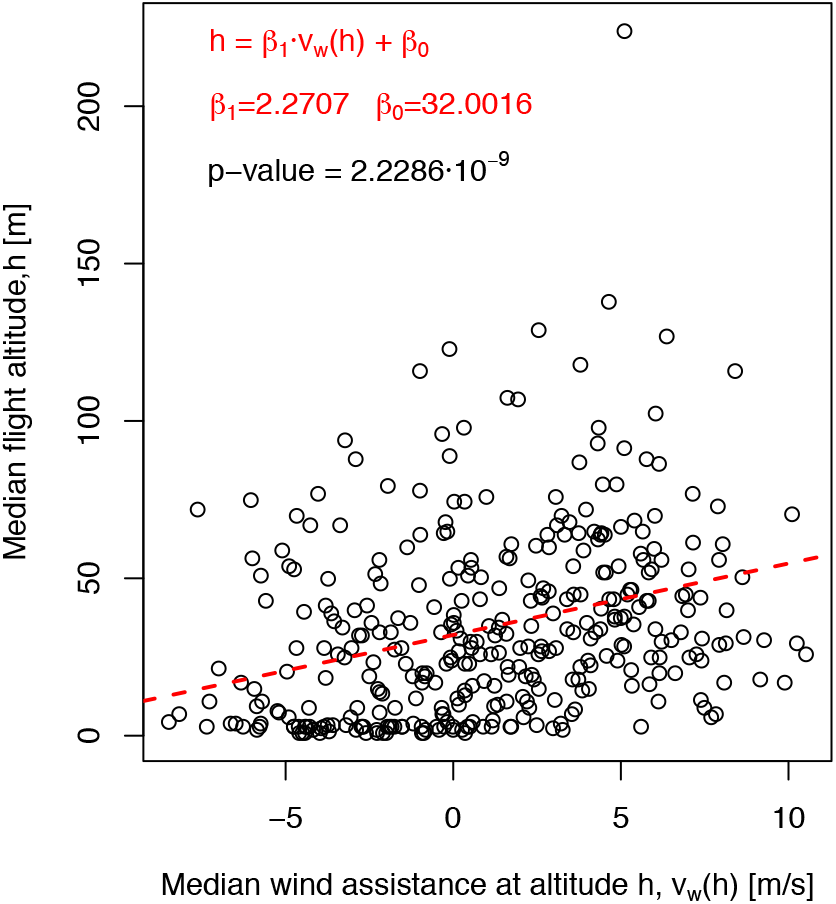
Gull’s median altitude *h* versus median wind assistance *V_w_* (head or tail wind) at median altitude *h* for 352 flights (16 distinct gulls). The regression line using *β*_1_ and *β*_0_ is plotted in red.

This statistical analysis tells us that gulls tend to maintain a ventral optic flow close to 25.23°/*s* whatever the wind conditions are while flying above the sea.

## 5. Takeoff time series analysis: individually tuned parametric model

In this section, 18 takeoffs are treated as independent observations despite these being recorded on 9 individual birds. Indeed, the weather, the wind, the state of the sea, the moment, and the fishing area were uncontrolled and different from one flight to another (Fig. 2b).

### (a) Parametric model estimation

The linear parametric models about each gull’s elevation dynamics were estimated with the System Identification Toolbox from the Matlab software (parameters: time constant *τ_h_* and static gain 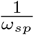 in (5.2). The maximum climbing speed *V_zmax_* (5.1) was computed from [24, 48]:

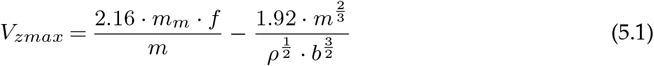

where *m_m_* is the mass of the flight muscles, *f* is the observed flapping frequency (3.26 Hz on average, see page 162 in [17]), *m* is the total mass including any added load, *ρ* is the air density (1.205 kg/m^3^ at 20° C) and *b* is the wing span. The vertical wind is low over the sea, consequently in flight, we neglected the vertical wind. For each of the 18 offshore takeoffs followed by a cruise flight, we took into account the morphological parameters of each gull.

### (b) Computation of the predicted altitude

The model output, i.e. the predicted altitude, *h_est_*, was computed with the Simulink environment from the Matlab software. The best fit factor of the optic flow-based control model is obtained by adjusting the flight muscle fraction 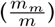 instead of the bird mass *m*, because the bird mass was known without any prey load. The fit factor considered was the goodness of fit between optimized simulated data (*h_est_*) and actual GPS data (*h_GPS_*) using a Matlab function with a normalized mean square error cost function (called NRMSE cost function). NRMSE fit factor varies between minus infinity (worse fit) to 1 (perfect fit). According to the table 15 in [21], the flight muscle ratio 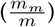 is relatively constant across birds species at 0.18 ± 0.05 (MEAN ± SD, with *n* = 221). Our simulated model has been adjusted with the flight muscle ratio in order to get the best fit factor, then adjusting the maximum climbing speed in the elevation dynamics model. For our group of 9 individual lesser black-backed gulls, we obtained the best fit factor with a corresponding distribution of flight muscle ratio 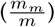 of 0.18 ± 0.03, which is quite similar to prediction 9 from [24]. The optic flow-based control model takes into account the observed correlation between the groundspeed *V_g_* and the altitude *h* coming from gulls’ GPS data. The proportionality factor is called here a ventral optic-flow set-point *ω_sp_* (2.6). Once the best fit factor has been reached by adjusting the flight muscle fraction 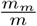, each gull’s altitude is re-computed by considering an altitude control model that directly feeds the elevation dynamics with a “target flight altitude”, noted an altitude set-point *h_sp_*, which is computed when the gull reached its steady-state altitude.

### (c) Optic flow-based altitude control model

We consider two scales of time. The gull’s forward dynamics responds faster than the gull’s upward dynamics (constrained by *V_zmax_* see (5.1)) because the height of flight arises from the response of a first order differential equation by considering the forward speed as a step input (5.2). The bird’s elevation dynamics is represented in Fig. 5a, this includes both the first order upward dynamics (5.2) and the maximum climbing speed *V_zmax_* (5.1).

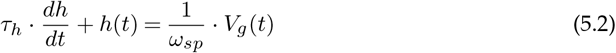

**Figure 5.**
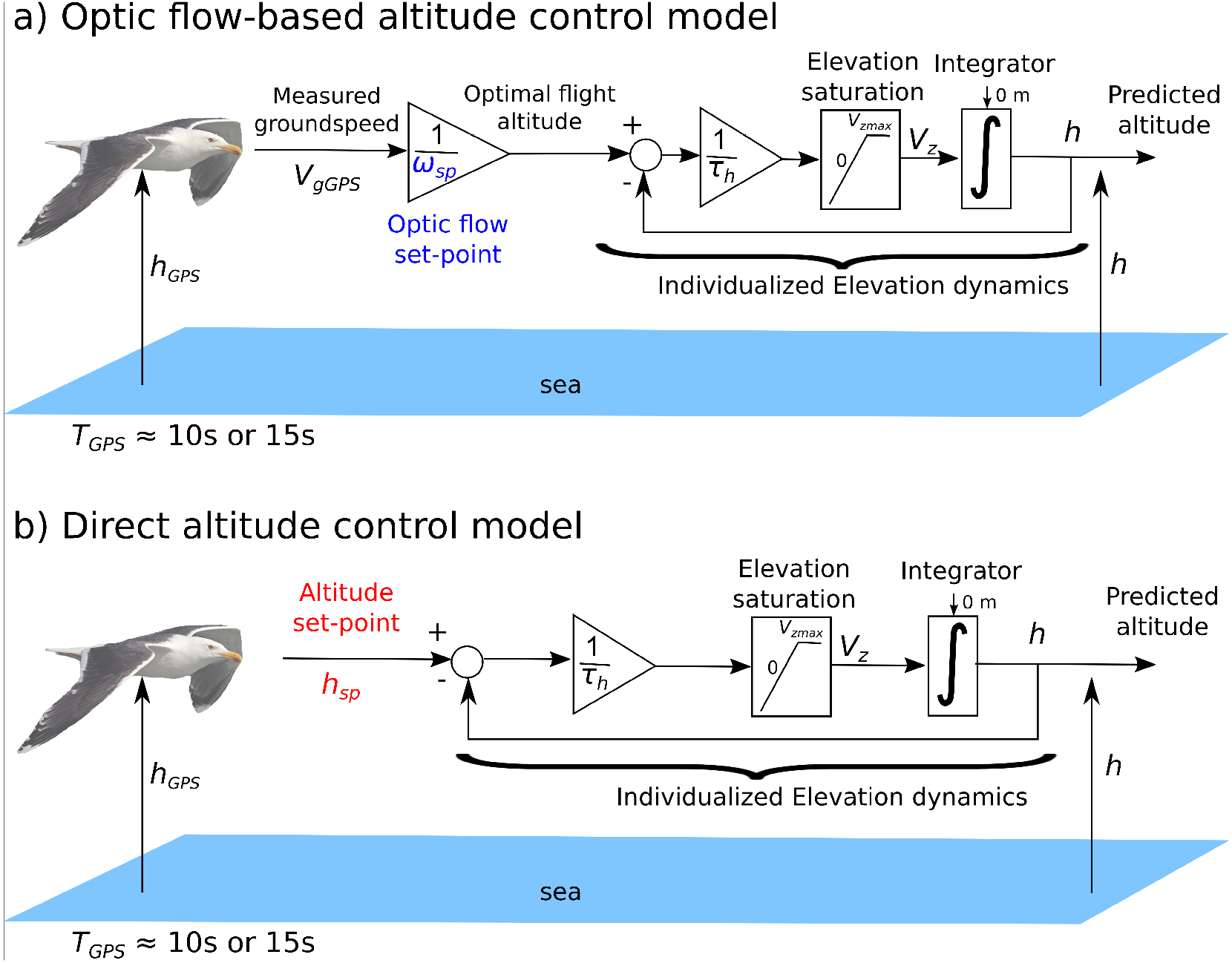
(a) Optic flow-based altitude control model including an individualized gull’s elevation dynamics. Once the gull has reached the minimum groundspeed to takeoff, groundspeed is then relatively constant during its flight, an optic-flow-based control system can be switched on and lead the gull to a given altitude depending on both its groundspeed *V_g_* and its ventral optic flow set-point *ω_sp_*. The ventral optic-flow set-point *ω_sp_* is an internal parameter used by the gull to tend asymptotically to its optimal flight altitude proportionally to its current groundspeed *V_g_*. This model correlates the gull’s current groundspeed to its current altitude. (b) Direct altitude control model. Here, the model only includes an individualized elevation dynamics and an altitude set-point *h_sp_*. This model does not impose asymptotically any proportionality between groundspeed and altitude. The altitude set-point *h_sp_* is an internal parameter used by the gull to select its “desired” or “target” flight altitude.

An explicit solution of equation (5.2) can be written, if we consider a step response at a given positive amplitude *V*_*g*0_ value, as follows:

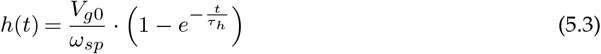

For each gull trajectory, we consider only one takeoff followed by a cruise flight, and then we perform a first order system identification described by the differential equation (5.2). In this model, a proportionality factor 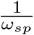 is introduced, which is the inverse of the ventral optic-flow set-point *ω_sp_* (2.6), and the input of the upward dynamics (5.2) is the groundspeed *V_g_*, which correlates the altitude *h* and the groundspeed *V_g_*. If the gull’s groundspeed is constant during takeoff as well as during cruising flight, then the predicted altitude profile will be the same with both models.

The inter-flight variability of the climb time constant (*τ_h_* = 97.3s ± 68.0s, with *n* = 18 takeoffs) was derived on the basis of morphological properties of the birds (*inter alia* age, wingspan, body mass including the load of prey and sex).

### (d) Direct altitude control model

Here, the bird’s elevation dynamics is represented in Fig. 5b, which includes both the first order upward dynamics (5.4) and the maximum climbing speed *V_zmax_* (5.1).

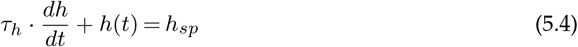

An explicit solution of equation (5.4) can be written, if we consider a step response at a given altitude *h_sp_* value, as follows:

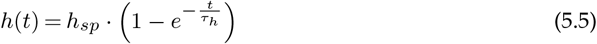

The “target flight altitude”, also called the altitude set-point is denoted *h_sp_*, which is computed from when the gull reached its steady-state altitude, i.e. the gull’s mean altitude when *t* > 3*τ_h_* or *t* > 5*τ_h_*, depending on data availability. In this model, there is no correlation between altitude and groundspeed.

### (e) Comparison between optic flow-based and direct altitude control models

A set of 18 trajectories representing 9 different gulls are individually shown in the horizontal plane in Fig. 2b. The set of GPS data are clustered and shown in Fig. 6 for the initial 400 seconds of each flight. It allows us not only to show the increase in speed during the gulls takeoff (Fig. 6a), but also their level flight along the vertical plane (Fig. 6b). Both groundspeed and altitude have been individually normalized by the steady state value reached by the gull’s groundspeed and altitude, respectively (Fig. 6). Consequently, both curves reach a steady state close to a value of one (Fig. 6).

**Figure 6.**
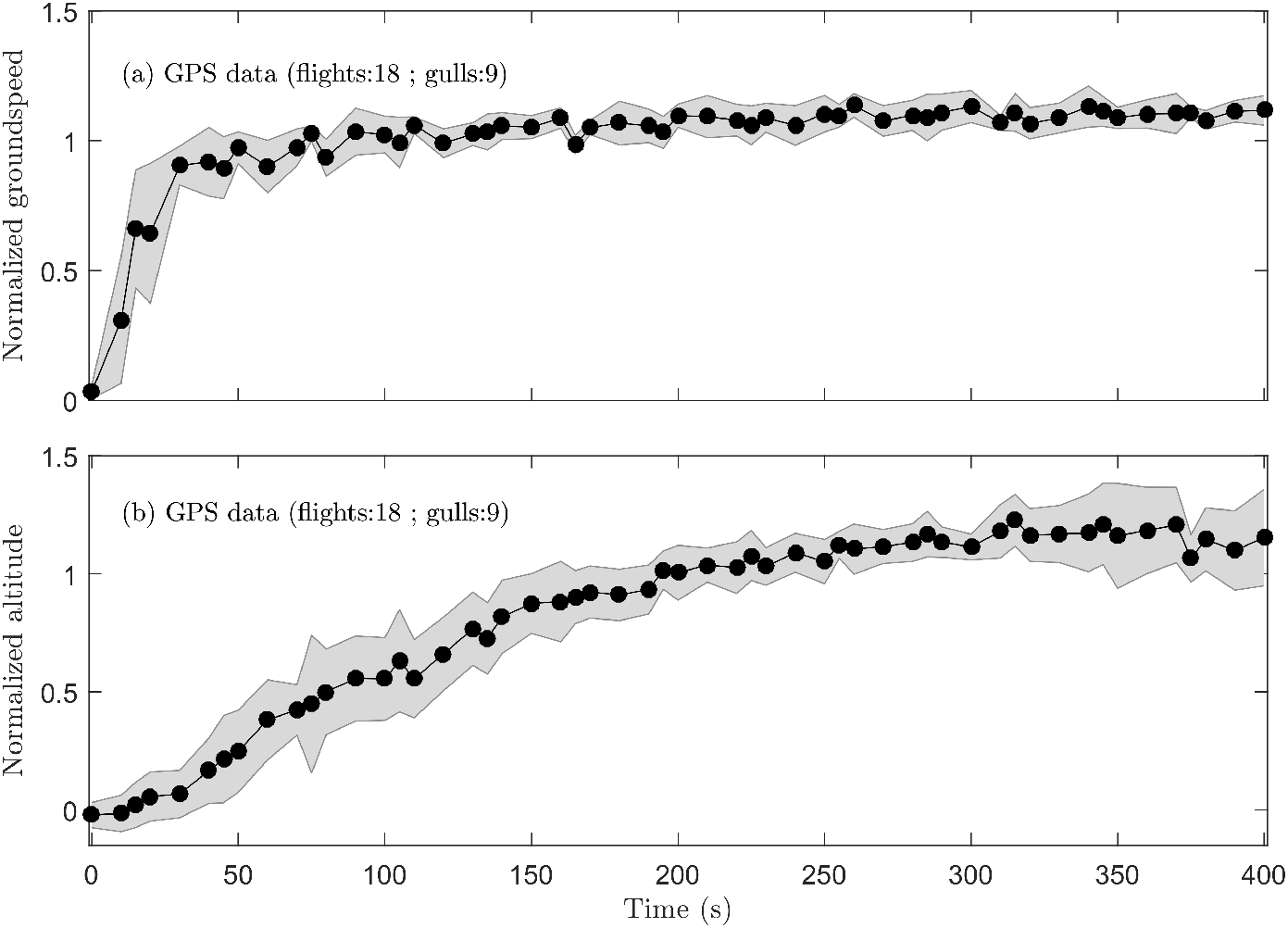
(a) Normalized groundspeed coming from GPS speed measurements 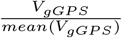, which is computed by the current groundspeed to average groundspeed ratio. (b) Normalized altitude coming from GPS data, which is computed by the current altitude to average altitude (by removing the first 100 seconds) ratio 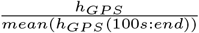. Black dots represent GPS data recorded at a sampling time 10 *s* (12 trajectories) or 15 *s* (6 trajectories). Each dot represents the median value and shaded areas represent the median absolute deviation (MAD) of the GPS data collected from 18 flights.

A linear 1st order parametric model on the data (18 trajectories) gives a fit factor value (i.e., a normalized mean square error cost function, called NRMSE cost function) of 40.4% on average (range: 10-80%). Then, by introducing a constraint on the climb rate according to prediction 10 in [24,48], a direct altitude control model based on a non-linear 1st order parametric model combined with an altitude set-point *h_sp_* (see Fig. 5b for details) gives a fit factor of on average 57.1% (range: 11-77%). However, by adding to the previous model a correlation between groundspeed and altitude, which is linked to what we call an optic flow set-point *ω_sp_* (see Fig. 5a for details), an optic flow-based control model gives a fit factor of 63.4% on average (range: 30-83%).

Examples comparing an optic flow-based control model to a direct altitude control model for one takeoff is given in Fig. 7b (the 17 other takeoffs are shown in Supplemental Information, Figs. S3-S19). We observe that in each case the fit factor was higher with an optic flow-based control model (blue dots in Fig. 7b rather than a direct altitude control model (red dots in Fig. 7b).

**Figure 7.**
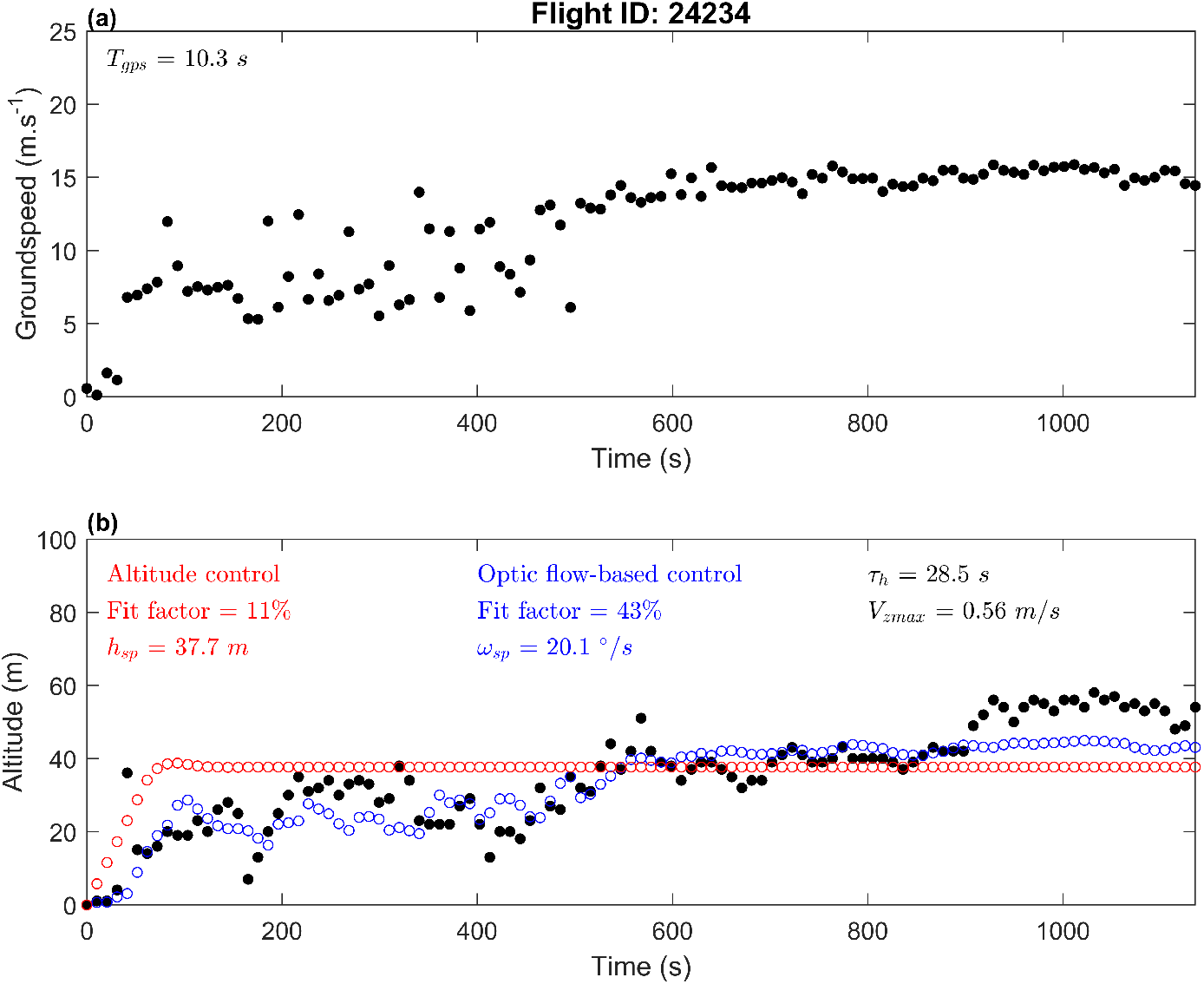
(a) Groundspeed of the gull ID24234 tracked with the GPS. (b) Altitude of the gull: black dots represent the GPS data, red dots represent the gull altitude on the basis of an altitude-based control model (fit factor: 11%), and the blue dots represent the gull altitude on the basis of an optic flow-based control model (fit factor: 43%). A significant correlation was observed between groundspeed and altitude of the GPS data (*ρ* =0.83, *p* ≪ 0.001 by Spearman’s test).

The set of normalized predicted altitudes (*n* = 18) computed with an altitude control model (Fig. 5b) is shown in Fig. 8a, and with an optic flow-based control model (Fig. 5b) is shown in Fig. 8b. Residuals, which are the errors between altitudes coming from GPS data and predicted altitudes coming from models, are represented in Figs. 8c-d. We compared the residuals distribution between the two models in transient response (white shaded boxes in Fig. S2) and in steady-state response (gray shaded boxes in Fig. S2). The median value of the residuals (Figs. 8c-d) coming from the optic flow-based model was significantly higher in transient response (one-sided Wilcoxon rank sum test, *n* = 27, *p* ≪ 0.001) and was also significantly higher in steady-state response (one-sided Wilcoxon rank sum test, *n* = 27, *p* ≪ 0.001). Consequently for both parts, the response predicted by the optic flow-based control model was better than the response predicted by the altitude control model. Finally, the average value of the residuals coming from each control model in transient response, then in steady-state response, were compared to a normal distribution centred around zero. The distributions of residuals with the optic flow-based control model (white shaded boxes in Fig. S2) were not significantly different from a normal distribution centred around zero (*t*-test, *n* = 27, *p* = 0.95 in transient response, and *p* = 0.07 in steady-state response). Residuals with the direct altitude control model (gray shaded boxes in Fig. S2) were significantly different from a normal distribution centred around zero (*t*-test, *n* = 27, *p* < 0.01 in transient response and *p* ≪ 0.001 in steady-state response). This statistical analysis shows that the optic flow-based control model is the most established model. Besides, for 13 out of 18 flights, we observe a significant correlation (Spearman’s test on GPS data) between groundspeed and altitude (*ρ* from 0.22 to 0.83, 13 flights). We therefore conclude that our optic flow-based control model (Fig. 5a) better explains the gulls’ GPS tracking data than the direct altitude control model (Fig. 5b).

**Figure 8.**
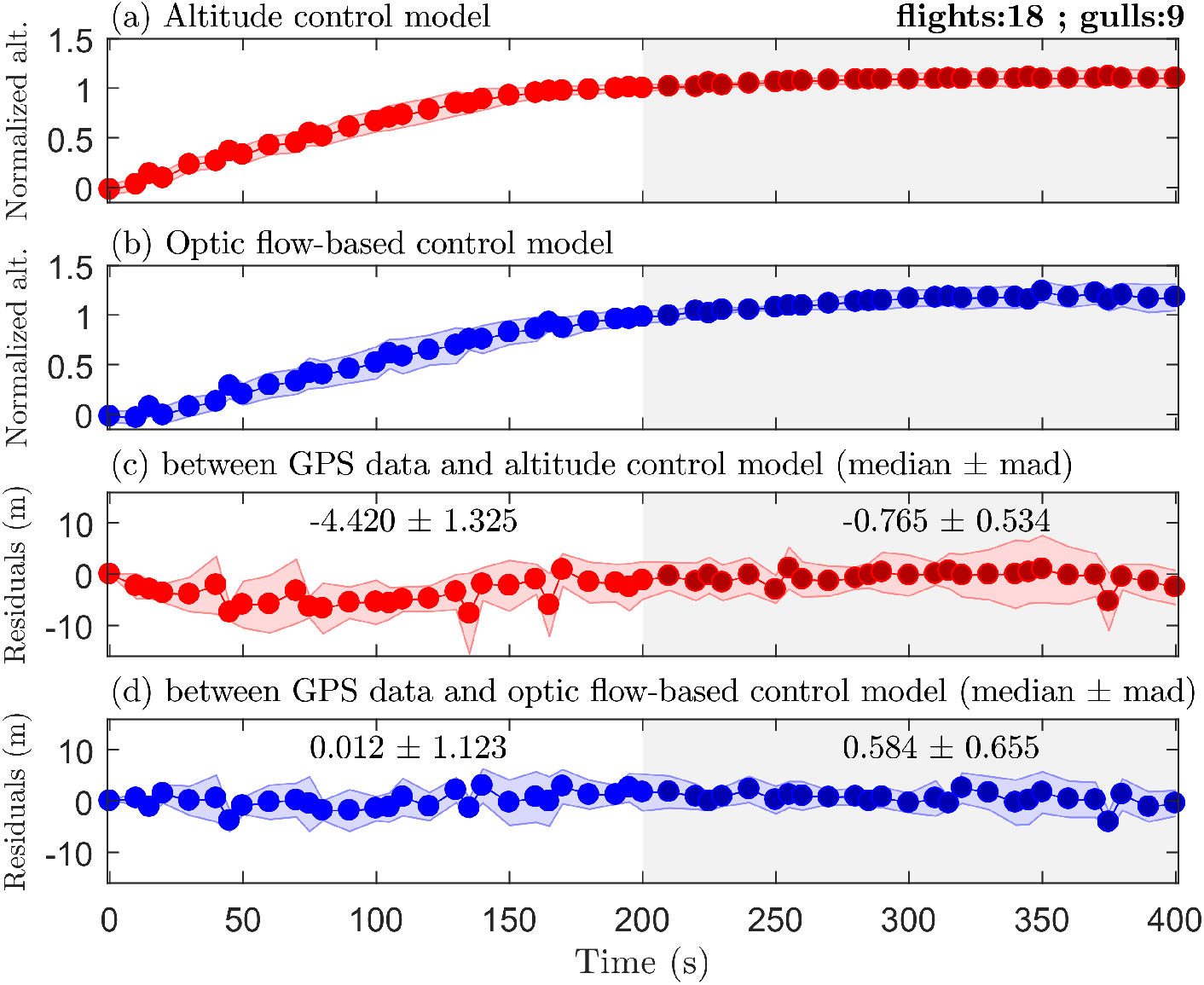
Red dots (altitude control model) or blue dots (optic flow-based control model) represent predicted altitude ((a) and (b)) or residuals ((c) and (d)) at a same sampling time 10 *s* (12 trajectories) or 15 *s* (6 trajectories) like GPS data (see Fig. 6). Each dot represents the median value and shaded areas represent the median absolute deviation (MAD) of data (*n* =18). The white shaded areas represent the transient response (time <200 *s*) during takeoff and ascent, and the gray shaded areas represent the steady state response (time >200 s) once in cruising flight. The duration 200 *s* ≈ 2 · *τ_h_* represents about 86% of the step response of a 1st order dynamic system (see (5.3)). (a) Normalized predicted altitude using an altitude control model (Fig. 5b), which is computed by current predicted altitude average predicted altitude (by removing the first 100 seconds) ratio 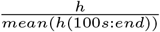. (b) Normalized predicted altitude using an **optic flow-based control model** (Fig. 5a), which is computed by current predicted altitude average predicted altitude (by removing the first 100 seconds) ratio 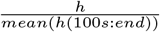. (c) Residuals between GPS data (Fig. 6b) and altitude computed with the altitude control model (data in (a)). (d) Residuals between GPS data (Fig. 6b) and altitude computed with the optic-flow based control model (data in (b)).

## 6. Discussion

### (a) Comparison of optic flow set-points identified by both analyses

We compared the distribution of ventral optic flow set-points coming from the tuned parametric model obtained from the takeoff time series (*ω_sp_* = 22°/*s* ± 9°/*s* with *n* = 18, Shapiro normality test: *p* = 0.16) and the parameter *ω_sp-lmer_* =25.23°/*s* obtained from the linear mixed effect model (4.2), respectively. No significant difference was observed between the *ω_sp_* distribution and the value *ω_sp-lmer_* (*t*-test, t:1.5296, df: 16, p-value:0.1457). This suggests shows that both analyses identify optic-flow set-points that are in the same range and not significantly different. As a consequence, both the takeoff time-series and the full dataset support the ventral optic flow regulation hypothesis in a consistent manner.

### (b) Effect of wind on the birds’ altitude

An additional outcome of the ventral optic flow regulation hypothesis [18,55] is that any increase in headwind will lead to a decrease in gull flight altitude in order to maintain the ventral optic flow constant (Fig. 9a). Conversely, any increase in tailwind will lead to an increase in bird altitude (Fig. 9c). A bird can adjust its ground speed by adjusting its airspeed or its heading relative to ground (and wind), thus allowing it to minimize its cost of transport in flight. The altitude control system based on optic flow is therefore consistent with previous observations on speed adjustment with respect to winds in migrating birds [2].

**Figure 9.**
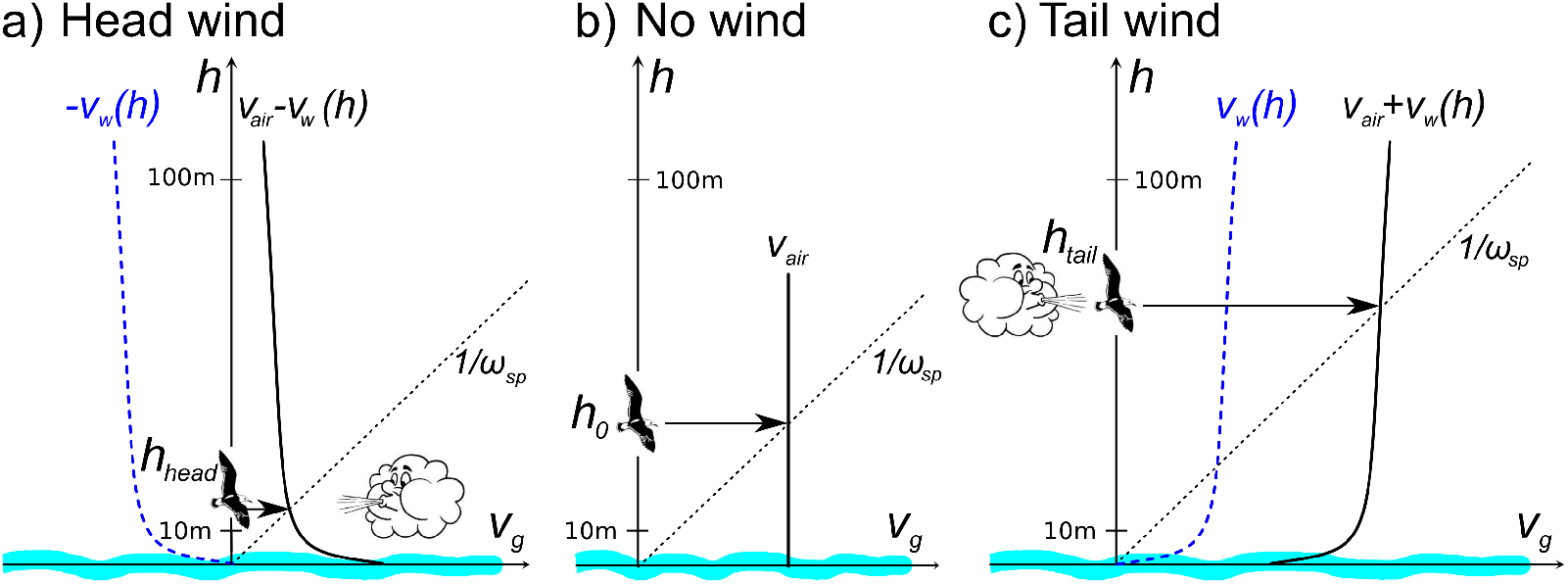
Gull’s speed and altitude for three different wind scenarios under the hypothesis that the gull adjusts its vertical lift to maintain constant its ventral optic flow. The red straight line 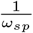 indicates the set of possible pairs of altitudes and groundspeeds allowed by the ventral optic flow regulation hypothesis. (a) In the presence of a head wind, given that the wind speed increases with the altitude, the groundspeed profile *V_air_* – *V_w_*(*h*) intersects the straight line 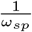 at a lower altitude *h_head_* than in absence of wind. (b) In the absence of wind, the ground speed and hence the altitude depend only on the airspeed produced by the agent: the vertical line *V_g_* = *V_air_* intersects the line 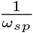 at the altitude *h*_0_. (c) In the presence of a tail wind, the ground speed profile *V_air_* + *V_w_*(*h*) intersects the straight line 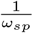 at a greater altitude *h_tail_* than in absence of wind. Modified from [55].

The small Hellman exponent α over relatively smooth surfaces, such as the sea, means that wind speed increases more rapidly than over a rough surface (e.g. a forest). Thus at higher altitudes (i.e., from 10 m to 100 m) wind speed will not vary much, but below 10 m wind speed can double going from 1 m to 10 m. Around the sea’s waves wind is deflected leading to a pattern of updrafts and downdrafts [50, 53, 68]. Together these effects are used by soaring seabirds in dynamic soaring, gust soaring or “sweeping flight” [50, 53, 68], and the characteristic meandering flight style that results has been termed “wave-meandering wing-sailing” [61]. Flapping seabirds can also use these features to gain a higher climb rate at the start of a take-off maneuver, taking off facing into the wind in the updraft formed by the deflection of the wind over a wave (see page 268 in [51], and [33]), which therefore reduces the effort required to take-off and accelerate to reach the maximum range speed *V_mr_*. Seabirds may also use the “ground effect” while flying very close to the sea to reduce their energetic expenditure [8], which is helpful for takeoff at sea.

### (c) Effect of altitude on optic flow

According to prediction 3 in [24, 48], the optimal altitude for a migratory bird is that where it can get just sufficient oxygen to maintain its cruising airspeed. This arises from the power required to fly at maximum range speed decreasing with altitude due to decreasing air density. Consequently, at an altitude of 6000 m, where the air density is half that at sea level, a bird should theoretically fly 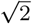 times faster. On the other hand, at a given optic-flow set-point working in a 100 m altitude range, the optic flow would be divided by a factor 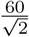 at an altitude of 6000 m. Therefore the optic flow would be too small to be maintained at the amplitude of the one generated in a 100 m altitude range. Recently, McLaren and colleagues (2016) analysing flights of lesser black-backed gulls flying between south-east England and The Netherlands recorded much greater flight altitudes than those observed here during homing flights to the breeding colony, with maximal values 1,240 m [42], even though typical values were lower at 100-150 m. On migratory flights, the gulls have been recorded flying higher still, though that is overland, with maximal altitudes around 5,000 m (unpublished data). Consequently, an optic flow based altitude control system can only work below a 100-meter altitude range where the optic flow is significant and detectable by the visual system of the birds.

### (d) Are groundspeed and altitude still proportional at higher altitudes?

Birds making lower altitude flights (<100-150 m) will generate a detectable optic flow. However, when on long distance or migratory flights birds may fly higher at hundreds to thousands of meters (see above), optic flow values will then be extremely low, thus unlikely to be suitable for regulating a given optic flow set-point. This relates to the finding for common swifts (*Apus apus*) by Hedenström & Åkesson (2017), that the swifts did not compensate for head- and tail winds as expected from flight mechanical theory when flying at high altitudes (>1000 m), but they did so at low altitudes (<100 m) [25]. This was interpreted as a failure to detect small changes in optic flow due to winds by the swifts’ visual system at high altitudes. In addition, for altitudes higher than 400 m, lesser black-backed gulls were observed to compensate less for cross-wind disturbance than they did at lower altitudes: fractional compensations were observed to decrease from about 1.3 (on average) to less than 0.5 at 900 m height [42]. At altitudes above 400 m, gulls’ groundspeed may be highly dependent on the wind speed: no altitude increase or decrease can be predicted with respect to the optic flow-based control model as optic flow is low thus its changes with altitude would be difficult to detect by the gulls’ visual system.

### (e) Bird navigation in the vertical dimension: can birds use barometric pressure to determine altitude?

The birds’ mechanoreceptive paratympanic organ (PTO) is located in the middle ear, and it is probably used by birds to detect barometric pressure [65]. Birds appear to use the PTO not only as a barometer to predict the onset of inclement weather [10,60,65], but also as a genuine altimeter to adjust their flight altitude during migration. Birds can fly level within ±20m for distances of 2-3km at altitudes of 700 – 1,100m, even at night [22], i.e. without visual cues. However, it is still an open question whether birds can use changing barometric pressure directly to measure their current altitude in real time.

A mechanoreceptive scale sensory organ found in fish [5] may play the same sensory function as the PTO in birds. It is known that fish can determine their depth using hydrostatic pressure [29, 63]. On this point, it was demonstrated that the dynamic depth sensing in fish is less than 1 m at a depth of 100 m [63]. However, water density is approximately 1,000 times higher than air density, and the pressure gradient in flight is therefore particularly low generating extremely low frequencies in the feedback signal to the bird’s elevation dynamics. Therefore, it would be difficult to adjust the flight altitude for a short period of time, only being practical for long periods of time such as for example during longer distance migratory flights.

### (f) Effect of wave motion on the optic flow pattern

The flight model assumes that the sea-surface, over which the gulls fly, provides a stationary reference frame: no data are currently available on the wave speed. Therefore, the optic flow experienced by the gulls is solely modeled as a function of their own movement (groundspeed and altitude). Previous studies on bird navigation over water suggest that the seascape (or more specifically the wavescape) is not a fixed reference frame [1], as the wave patterns move, usually in roughly the same direction as the wind but at a slower speed. Therefore the perceived optic flow will be different than the physical optic flow. Alerstam & Petterson (1976) suggested that the motion of the wave scape allows birds to only partially compensate for wind-drift over the sea [3], thus presumably a similar constraint may apply to using the ventral optic flow for control of flight altitude.

Overall, the wave pattern will reduce the adjustment of altitude if a fixed optic flow set-point was used, as under headwinds perceived optic flow will be higher than otherwise, i.e. even as groundspeed approaches zero there will still be a perceived optic flow if the wavescape is moving, which would lead to higher flight altitudes than expected. While under tailwinds optic flow is somewhat reduced, as the sea surface pattern will be moving in the same direction as the bird, and hence lower than expected flight altitudes would result. The wave pattern distorts the ventral optic flow perceived: such disturbances could be added to the flight model once data or a methodology of how to obtain wave pattern becomes available.

However, for optic flow to be useful ripples above the sea are essential to form a textured surface. In fact, it was observed by Heran & Lindauer (1963) that a great number of honeybees plunged into the water when the water surface was mirror smooth [28]. An altitude control system based solely on a ventral optic flow regulation irrevocably pulls any flying animal down whenever its eye fails to measure an optic flow [18]. This did not happen in honeybees when the water surface was rippled [28, 62] or when a floating bridge provided a visual contrast [28].

At this level of reasoning, we may wonder if the visual pattern produced by waves was textured enough during the gulls’ flights for an optic flow field to be perceived. To investigate this, knowing that the average significant wave height of the Baltic Sea in 1991–2015 was in the range 0.44–1.94 m [34], which corresponds to a Beaufort number of 3 (gentle breeze, mean wind speed equivalent from 3.4 m/s to 5.4 m/s) to 4 (moderate breeze, mean wind speed equivalent from 5.5 m/s to 7.9 m/s) [6]. We deduce that gulls could see scattered or fairly frequent white-crested waves at an effective height of 10 m above the sea level. However for Beaufort numbers from 0 to 2, the sea has a smooth appearance, which makes for poor visual conditions to perceive an optic flow field. Interestingly, the wind conditions corresponding to a Beaufort number from 3 to 4 fit not only with the wind conditions of gulls in flight (Fig. S20), but also with their altitude (see page 166 in [17]). We can conclude that wind is an important parameter to generate an optic flow field cue, and to help gulls to control their flight above the sea.

Little is known about the visual system of gulls. The spatial acuity of seabirds can be more than four times lower than that in humans [43], with a maximum spatial acuity of about 60 cycles/degree in humans. Moreover, in seabirds rods are evenly distributed across the entire retina [15], which allow them to conveniently detect the optic flow coming from the sea. Most of the seabirds have a maximum binocular field width in the 15° – 30° range (about 120° in humans), which is limited, suggesting that binocular vision plays only a minor role in seabirds’ flight control system [39]. We conclude that the optic flow field is the major visual cue used by seabirds to control their flight above the sea.

### (g) Optic-flow set-point: differences between honeybees and gulls

There are a number of differences in flight behaviours expressed by birds and flying insects [4]. Typically, the average maximum airspeed of honeybees is approximately 7.5 m/s with a minimum power speed of their power U-curve at 3.3 m/s [44]. In free-flight natural conditions, honeybees have been observed to fly from 3.3 m/s to 5.1 m/s [44]. However, lesser black-backed gulls typically fly at an airspeed in natural offshore conditions at an average 12.3 m/s ± 2 m/s (see [17], page 166) with a minimum power speed of their power U-curve at 9.3 m/s (computed for lesser black-backed gull, see [27]). Hence, lesser black-backed gulls can fly 3 times faster than honeybees by comparing their minimum power speed.

In honeybees, average maximal flight height is about 2.5 m over natural terrain [16, 28]. In general, lesser black-backed gulls fly at an altitude over sea of up to 130 m with a distribution of 31 m ± 29 m on average (see [17], pages 166-167) during foraging flights. We conclude that lesser black-backed gulls fly much higher than honeybees during foraging flights, which reduces optic flow emanating from the sea.

Consequently, we can conclude from these two last points that the ventral optic-flow set-point of lesser black-backed gulls is much lower than that typically experienced by honeybees, knowing that the ventral optic-flow set-point of honeybees is close to 200°/*s*. Our statistical analysis estimates that the ventral optic-flow set-point of lesser black-backed gulls is close to 25°/*s* on average (see section 4), which is a detectable value by the gulls’ visual system [39,40, 41]. A recent review indicates that pigeons’ fast LM neurons (pretectal nucleus lentiformis mesencephali) respond to optic flow stimuli of their preferred backward direction (front to back visual stimuli: temporal to nasal on the retina) in this same angular velocities range [69].

## 7. Conclusion

A mathematical model of optic flow-based offshore takeoff control system in lesser black-backed gulls was developed in this study to understand what visual cue can be used by seabirds to control their takeoff and to cruise over a sea surface. This mathematical model introduced an optic flow set-point parameter, which aims to be maintained constant by seabirds during take-off manoeuvers and cruising foraging flights. Besides, the model takes into account the bird’s individual morphology through its elevation dynamics. Finally, both analyses on the takeoff time-series and the full dataset support the ventral optic flow regulation hypothesis in a consistent manner.

We conclude that the optic-flow regulation principle allows seabirds to control their altitude over sea at low flight altitudes without having to measure their current altitude directly by another method. To do this, they just have to measure the optic flow perceived from the sea to adjust their vertical thrust in order to maintain the ventral optic flow at a given value, called the optic-flow set-point, as previously suggested for flying insects [18, 55]. According to both the airspeed and altitude ranges of lesser black-backed gulls measured during flight in their natural environment, we demonstrate that gulls could control their altitude by regulating the ventral optic-flow at a value of 25°/*s* on average, allowing them to fly jointly up to 130 m in altitude at a groundspeed up to 20 *m/s*, while maintaining visual contact with the sea. The introduction of this asymptotic proportionality relationship for birds also accounts very nicely for the transient altitude response during takeoff. Overall, gulls need such accurate altitude control based on optic flow to optimize their energetic effort irrespectively of favorable or unfavorable unknown wind conditions while being robust to ground disturbances such as relief.

## Supporting information

Data set of full flights - statistical analysis

data set of gulls takeoffs - Flight modelling

## Authors’ contributions

JRS, FR, TJE and AH developed the modelling; JRS ran the models on Matlab software; TJE and SÅ tagged the gulls and collected the data; JRS, TJE and FR analysed the modelling results; JSB provided the tracking system; FR and AH supervised the collaboration; FR drew figs 1 and 9; TJE drew figs 2 and 3; JRS drew figs 5, 6, 7, 8 and S1-S19; TJE, JRS and FR drew fig 4; JRS wrote the first draft of the paper; all authors prepared and revised the manuscript.

## Acknowledgement

French financial support was provided by a project grant to FR from the CNRS PEPS Avimod (Exomod Program). FR and JRS were also supported by Aix Marseille Université and CNRS (Life Science; Information Science, and Engineering Science and Technology). Financial support was via project grants to SÅ and AH from the Swedish Research Council (621-2007-5930; 621-2010-5584, 621-2013-4361, 621-2012-3585) and Lund University, and a Linnaeus grant from the Swedish Research Council (349-2007-8690) to the Centre for Animal Movement Research (CAnMove) and Lund University. Field work was partly supported by WWF Sweden. UvA-BiTS is facilitated by infrastructures for eScience, developed with support of the NLeSC (http://www.esciencecenter.com/) and LifeWatch, carried out on the Dutch national e-infrastructure with support of SURF Foundation. Permissions to capture and ring birds were from the Swedish Nature Protection Board (Naturvårdsverket) and the Swedish Ringing Office at the Natural History Museum in Stockholm. Ethical permission to tag the gulls was from Malmö/Lund Djurförsöksetiska nämnd (No. M112-09, M470-12). Permission to work in the protected area was from the county administration Länstyrelsen Gotlands Län. For assistance during fieldwork we thank the whole Baltic Seabird Group team, especially: Olof Olsson, Jonas Hentati-Sundberg, Per-Arvid Berglund, Aron Hejdström, Martina Kadin, Natalie Isaksson, and Rebecca Young. We would like to thank Prof. Maan El Badaoui El Najjar (Univ. of Lille) for his fruitful comments during this analysis.

