## Supplementary material for "Optic flow helps explain gulls’ altitude control over seas": Data set of full flights - statistical analysis: full_flights_analysis.R

```
# Introduction ----
#' This script runs the analysis presented in the Full flights’
#' dataset analysis presented in Serres et al. 2019. Optic flow
#' cues help explain altitude control over sea in freely flying
#' gulls. J. R. Soc. Interface


# Load required packages ----
# Install required R packages
install.packages(c("lme4", "dplyr", "pbkrtest"))

# Load packages ----
library(lme4)
library(dplyr)
library(pbkrtest)


# Set working directory ----
# If needed set working directory (this should be where the data file is located)
# See help for how to use
?setwd()

# Load flight dataset  ----
# Load in the data (assumes it is in your working directory)
flights_sub <- read.csv("flights_summary.csv")


# Filter flight data -----
# Exclude flights where median altitude is <= 0 m
flights_over0 <- filter(flights_sub, altitude_callib > 0)

# Recalculate wind speed for at flight height -----
#windspeed_corrected(k)=a(k,9).*(a(k,5) ./ href).^alpha
alpha<-0.11
href<-10

flights_over0$wind_flight_height <- flights_over0$track_head_wind_10m*(flights_over0$altitude_callib / href)^alpha

# LMM model -----
# Run model. The second part +(1|ring_number) is the random part, this allows for random intercepts for the different birds. The slopes will still be the same.
mod05 <- lmer((altitude_callib) ~
                wind_flight_height +
                (1|ring_number),
              data = flights_over0)

# View model output
mod05
summary(mod05)
anova(mod05)


# Calculate statistical significance ----
# Significance test function
KRSumFun <- function(object, objectDrop, ...) {
  krnames <- c("ndf","ddf","Fstat","p.value","F.scaling")
  r <- if (missing(objectDrop)) {
    setNames(rep(NA,length(krnames)),krnames)
  } else {
    krtest <- KRmodcomp(object,objectDrop)
    unlist(krtest$stats[krnames])
  }
  attr(r,"method") <- c("Kenward-Roger via pbkrtest package")
  r
}

# Calculate significance
# P values for altitude vs head wind speed
drop1(mod05, test="user", sumFun=KRSumFun)

# Plot data and model fit ----
plot(altitude_callib ~
       wind_flight_height,
     data = flights_over0, xlab=expression('Median wind assistance at altitude h, v'  [w] * '(h) [m/s]'), ylab="Median flight altitude,h [m]")
# Show model line
abline(a = 32.0016, b = 2.2707, lty = 2, lwd = 2, col = "red")
text(-4, 220, expression('h = ' * beta [1] * '·v'  [w] * '(h) + ' * beta [0]) , col="red") 
text(-2.45, 200, expression(beta [1]  *'=2.2707   '  *  beta [0] *'=32.0016'), col="red")
text(-2.82, 180, expression('p-value = 2.2286·10'  ^-9))
```
