## Supplementary material for "Optic flow helps explain gulls’ altitude control over seas": Data set of full flights - statistical analysis: full_flights_analysis_file_description.txt

Data and R code required to run the analysis presented in Figure 6 and section 4(a) of Serres et al. 2019. Optic flow cues help explain altitude control over sea in freely flying gulls. J. R. Soc. Interface

flight_summary.csv
Data file containing details on flights tracked of lesser black-backed gulls.
flight_id_combined - ID of flight tracked
device_info_serial - ID of GPS tracking device
ring_number - Bird identification ring number
n - Number of GPS locatiosn tracked during flight
altitude_callib - mean flight altitude during flight
ecmwf_wind_10m_dir - mean wind direction (10 m altitude)
ecmwf_wind_10m_speed - mean wind speed (10 m altitude)
vg - ground speed
track_head_wind_10m - wind speed in track direction
track_cross_wind_10m - wind speed perpendicular to track direction


full_flights_analysis.R
R script containing analysis code used.
R code is commented to provide explanation of steps taken.
