## Supplementary material for "Optic flow helps explain gulls’ altitude control over seas": data set of gulls takeoffs - Flight modelling: Data_ID436.m

```
%% Data flight Gull ID436 
%%
clear all
clc
close all

ID_gull = 8114315 ;
ID_flight = 436;
sex = 'M';
t= sprintf('Flight ID: %d', ID_flight);

altitude1 = importdata('C:\Users\VODKA\Desktop\Birds Flight\Data Birds\altitudeID436.txt');
altitude = altitude1(1:end-7);
speed1 = importdata('C:\Users\VODKA\Desktop\Birds Flight\Data Birds\speedID436.txt');
speed = speed1(1:end-7);
Te = 1080/76 ; % GPS sampling rate
time  = (0:1:length(altitude)-1)*Te; % 
tmax = max(time)-7*Te; % aucune suppression de point
g = 9.82; % Swedish gravity (m/s)
Tau = 138.33; % time constant of the elevation dynamics (s)
K = 3.29; % static gain of the elevation dynamics (SI)
altitude_sp = mean(altitude(round(5*Tau/Te):floor((tmax-0*Te)/Te)));

m = 0.780; % bird mass en kg
f = 3.26; % Wing beat frequency en Hz
fmf = 0.19; % flight muscle fraction
% cite: A. Hedenstöm (2003) Twenty-three testable predictions about bird
% flight, and C.J. Pennycuick (2008) Modelling the flying bird (Academic Press)

rho20 = 1.204; % air volomical mass at 20°C (kg/m^3)
b = 1.518; % wingspan en m

Vzmax = (2.16*fmf*f)-1.92*m^(2/3)/(rho20^(1/2)*b^(3/2)); % Max elevation speed (m/s)
% cite: A. Hedenstöm (2003) Twenty-three testable predictions about bird
% flight,


for n = 1:(length(altitude)-1)
        Vz(n) = (altitude(n+1)-altitude(n))/Te;
end

time_436 = time/Tau ;
altitude_436 = altitude/max(altitude);
%return

%% Import U-curve from Tom Evans
% Computed with: mass m = 0.742kg ; wingspan b = 1.4m ; wing area S = 0.177m²
% ; wingbeat frequency f = 3.26Hz
%% Import data from text file.
% Script for importing data from the following text file:
%
%    C:\Users\VODKA\Desktop\Gulls_mathieu_avizou\flight_power.csv
%
% To extend the code to different selected data or a different text file,
% generate a function instead of a script.

% Auto-generated by MATLAB on 2017/01/17 17:49:56

%% Initialize variables.
filename = 'flight_power.csv';
delimiter = ',';
startRow = 2;

%% Format string for each line of text:
%   column2: double (%f)
%	column3: double (%f)
% For more information, see the TEXTSCAN documentation.
formatSpec = '%*s%f%f%*s%*s%*s%*s%*s%*s%*s%*s%*s%*s%*s%*s%*s%*s%*s%*s%*s%*s%*s%*s%*s%[^\n\r]';

%% Import data from text file.
% Script for importing data from the following text file:
%
%    /Users/mathieu/Documents/MATLAB/Gulls_mathieu_avizou/flight_power.csv
%
% To extend the code to different selected data or a different text file,
% generate a function instead of a script.

% Auto-generated by MATLAB on 2017/01/19 18:02:04

%% Initialize variables.
filename = 'flight_power.csv';
delimiter = ',';
startRow = 2;

%% Format for each line of text:
%   column2: double (%f)
%	column3: double (%f)
% For more information, see the TEXTSCAN documentation.
formatSpec = '%*s%f%f%*s%*s%*s%*s%*s%*s%*s%*s%*s%*s%*s%*s%*s%*s%*s%*s%*s%*s%*s%*s%*s%[^\n\r]';

%% Open the text file.
fileID = fopen(filename,'r');

%% Read columns of data according to the format.
% This call is based on the structure of the file used to generate this
% code. If an error occurs for a different file, try regenerating the code
% from the Import Tool.
dataArray = textscan(fileID, formatSpec, 'Delimiter', delimiter, 'HeaderLines' ,startRow-1, 'ReturnOnError', false, 'EndOfLine', '\r\n');

%% Close the text file.
fclose(fileID);

%% Post processing for unimportable data.
% No unimportable data rules were applied during the import, so no post
% processing code is included. To generate code which works for
% unimportable data, select unimportable cells in a file and regenerate the
% script.

%% Allocate imported array to column variable names
airspeed = dataArray{:, 1};
mechanical_power = dataArray{:, 2};


%% Clear temporary variables
clearvars filename delimiter startRow formatSpec fileID dataArray ans;
```
