## Supplementary material for "Optic flow helps explain gulls’ altitude control over seas": data set of gulls takeoffs - Flight modelling: multi_graph2.m

```
clear all
clc
close all

%fitfactor = [66 80 71 40 17 65 40 59 71 70 40 80 69 61 15 63 59 62];
%omega_sp = [17 14 21 35 40 22 20 11 16 9 20 14 26 23 45 35 16 27];
%tau = [138 104 101 28 38 37 44 54 218 261 29 55 131 182 57 125 37 57];

% Shaded nomalized graphs
% Data at Te = 15s
N15 = 27 ; % number of points
alt15 = [];
spd15 = [];
Time15 = 0:15:(N15-1)*15;
%Time = 0:10:(N10-1)*10;
Time10 = [10:30:400 20:30:400];

t = [Time15 Time10];
[t_sort,t_index] = sort(t);
NaN15 = NaN(length(Time10),1); 

% Useful to get residuals in meters
% Factor in order to denormalize
residuals_factor_gps = [];
% Useful  to compute residuals in meters of the OF-based altitude control
% Factor in order to denormalize
residuals_factor_est = [];
% Useful to compute residuals in meters od the direct altitude control
% Factor in order to denormalize
residuals_factor_est_direct_alt = [];

load('FlightID436.mat')
Te = 1080/76 ; % 14.21 s OK
alt15 = [altitude(1:N15)/mean(altitude(7:N15)) ; NaN15];
est_alt15 = [est_altitude(1:N15)/mean(est_altitude(7:N15)) ; NaN15];
spd15 = [speed(1:N15)/mean(speed(1:N15)) ; NaN15];

residuals_factor_gps = [residuals_factor_gps mean(altitude(7:N15)) ];
residuals_factor_est = [residuals_factor_est mean(est_altitude(7:N15))];

% GPS altitude
alt15 = alt15(t_index);
% Altitude estimated on the basis of ground speed and optic flow set-point
est_alt15 = est_alt15(t_index);
% GPS groundspeed
spd15 = spd15(t_index);

clear altitude est_altitude speed

load('FlightID564.mat')
Te = 1755/118 ; % 14.87 s OK
alt15_add = [altitude(1:N15)/mean(altitude(7:N15)); NaN15];
alt15_add(1:end) = alt15_add(t_index);
alt15 = [alt15 alt15_add];
est_alt15_add = [est_altitude(1:N15)/mean(est_altitude(7:N15)) ; NaN15];
est_alt15_add = est_alt15_add(t_index);
est_alt15 = [est_alt15 est_alt15_add];
spd15_add =[speed(1:N15)/mean(speed(1:N15)) ; NaN15] ;
spd15_add = spd15_add(t_index);
spd15 = [spd15 spd15_add];

residuals_factor_gps = [residuals_factor_gps mean(altitude(7:N15)) ];
residuals_factor_est = [residuals_factor_est mean(est_altitude(7:N15))];

clear alt15_add est_alt15_add spd15_add
clear altitude est_altitude speed

load('FlightID571.mat')
Te = 915/59; % 15.5 s OK
alt15_add = [altitude(1:N15)/mean(altitude(7:N15)); NaN15];
alt15_add(1:end) = alt15_add(t_index);
alt15 = [alt15 alt15_add];
est_alt15_add = [est_altitude(1:N15)/mean(est_altitude(7:N15)) ; NaN15];
est_alt15_add = est_alt15_add(t_index);
est_alt15 = [est_alt15 est_alt15_add];
spd15_add =[speed(1:N15)/mean(speed(1:N15)) ; NaN15] ;
spd15_add = spd15_add(t_index);
spd15 = [spd15 spd15_add];

residuals_factor_gps = [residuals_factor_gps mean(altitude(7:N15)) ];
residuals_factor_est = [residuals_factor_est mean(est_altitude(7:N15))];

clear alt15_add est_alt15_add spd15_add
clear altitude est_altitude speed

load('FlightID5917.mat')
Te = 1140/77 ; % 14.80 s OK
alt15_add = [altitude(1:N15)/mean(altitude(7:N15)); NaN15];
alt15_add(1:end) = alt15_add(t_index);
alt15 = [alt15 alt15_add];
est_alt15_add = [est_altitude(1:N15)/mean(est_altitude(7:N15)) ; NaN15];
est_alt15_add = est_alt15_add(t_index);
est_alt15 = [est_alt15 est_alt15_add];
spd15_add =[speed(1:N15)/mean(speed(1:N15)) ; NaN15] ;
spd15_add = spd15_add(t_index);
spd15 = [spd15 spd15_add];

residuals_factor_gps = [residuals_factor_gps mean(altitude(7:N15)) ];
residuals_factor_est = [residuals_factor_est mean(est_altitude(7:N15))];

clear alt15_add est_alt15_add spd15_add
clear altitude est_altitude speed

load('FlightID14477.mat')
Te = 1892/120 ; % 15.76 s OK
alt15_add = [altitude(1:N15)/mean(altitude(7:N15)); NaN15];
alt15_add(1:end) = alt15_add(t_index);
alt15 = [alt15 alt15_add];
est_alt15_add = [est_altitude(1:N15)/mean(est_altitude(7:N15)) ; NaN15];
est_alt15_add = est_alt15_add(t_index);
est_alt15 = [est_alt15 est_alt15_add];
spd15_add =[speed(1:N15)/mean(speed(1:N15)) ; NaN15] ;
spd15_add = spd15_add(t_index);
spd15 = [spd15 spd15_add];

residuals_factor_gps = [residuals_factor_gps mean(altitude(7:N15)) ];
residuals_factor_est = [residuals_factor_est mean(est_altitude(7:N15))];

clear alt15_add est_alt15_add spd15_add
clear altitude est_altitude speed

load('FlightID16542.mat')
Te = 1327/83 ; % 15.98 s OK
alt15_add = [altitude(1:N15)/mean(altitude(7:N15)); NaN15];
alt15_add(1:end) = alt15_add(t_index);
alt15 = [alt15 alt15_add];
est_alt15_add = [est_altitude(1:N15)/mean(est_altitude(7:N15)) ; NaN15];
est_alt15_add = est_alt15_add(t_index);
est_alt15 = [est_alt15 est_alt15_add];
spd15_add =[speed(1:N15)/mean(speed(1:N15)) ; NaN15] ;
spd15_add = spd15_add(t_index);
spd15 = [spd15 spd15_add];

residuals_factor_gps = [residuals_factor_gps mean(altitude(7:N15)) ];
residuals_factor_est = [residuals_factor_est mean(est_altitude(7:N15))];

clear alt15_add est_alt15_add spd15_add
clear altitude est_altitude speed
% Data at Te = 10s
% N = 41 ; % number of points
% alt = [];
% est_alt10 = [];
% spd = [];
clear t t_sort t_index

N10 = 41;
Time15 = 15:30:400;
Time10 = 0:10:(N10-1)*10;

t = [Time10 Time15];
[t_sort,t_index] = sort(t);
NaN10 = NaN(length(Time15),1); 

load('FlightID5225.mat')
Te = 810/82 ; % 9.87s
alt15_add = [altitude(1:N10)/mean(altitude(10:N10)); NaN10];
alt15_add(1:end) = alt15_add(t_index);
alt15 = [alt15 alt15_add];
est_alt15_add = [est_altitude(1:N10)/mean(est_altitude(10:N10)) ; NaN10];
est_alt15_add = est_alt15_add(t_index);
est_alt15 = [est_alt15 est_alt15_add];
spd15_add =[speed(1:N10)/mean(speed(1:N10)) ; NaN10] ;
spd15_add = spd15_add(t_index);
spd15 = [spd15 spd15_add];

residuals_factor_gps = [residuals_factor_gps mean(altitude(10:N10)) ];
residuals_factor_est = [residuals_factor_est mean(est_altitude(10:N10))];

clear alt15_add est_alt15_add spd15_add
clear altitude est_altitude speed

load('FlightID24611.mat')
Te = 10.29 ; 
alt15_add = [altitude(1:N10)/mean(altitude(10:N10)); NaN10];
alt15_add(1:end) = alt15_add(t_index);
alt15 = [alt15 alt15_add];
est_alt15_add = [est_altitude(1:N10)/mean(est_altitude(10:N10)) ; NaN10];
est_alt15_add = est_alt15_add(t_index);
est_alt15 = [est_alt15 est_alt15_add];
spd15_add =[speed(1:N10)/mean(speed(1:N10)) ; NaN10] ;
spd15_add = spd15_add(t_index);
spd15 = [spd15 spd15_add];

residuals_factor_gps = [residuals_factor_gps mean(altitude(10:N10)) ];
residuals_factor_est = [residuals_factor_est mean(est_altitude(10:N10))];

clear alt15_add est_alt15_add spd15_add
clear altitude est_altitude speed

load('FlightID16606.mat')
Te = 887/85 ; % 10.4 s
alt15_add = [altitude(1:N10)/mean(altitude(10:N10)); NaN10];
alt15_add(1:end) = alt15_add(t_index);
alt15 = [alt15 alt15_add];
est_alt15_add = [est_altitude(1:N10)/mean(est_altitude(10:N10)) ; NaN10];
est_alt15_add = est_alt15_add(t_index);
est_alt15 = [est_alt15 est_alt15_add];
spd15_add =[speed(1:N10)/mean(speed(1:N10)) ; NaN10] ;
spd15_add = spd15_add(t_index);
spd15 = [spd15 spd15_add];

residuals_factor_gps = [residuals_factor_gps mean(altitude(10:N10)) ];
residuals_factor_est = [residuals_factor_est mean(est_altitude(10:N10))];

clear alt15_add est_alt15_add spd15_add
clear altitude est_altitude speed

load('FlightID20104.mat')
Te = 2165 /208 ; % 10.4 s
alt15_add = [altitude(1:N10)/mean(altitude(10:N10)); NaN10];
alt15_add(1:end) = alt15_add(t_index);
alt15 = [alt15 alt15_add];
est_alt15_add = [est_altitude(1:N10)/mean(est_altitude(10:N10)) ; NaN10];
est_alt15_add = est_alt15_add(t_index);
est_alt15 = [est_alt15 est_alt15_add];
spd15_add =[speed(1:N10)/mean(speed(1:N10)) ; NaN10] ;
spd15_add = spd15_add(t_index);
spd15 = [spd15 spd15_add];

residuals_factor_gps = [residuals_factor_gps mean(altitude(10:N10)) ];
residuals_factor_est = [residuals_factor_est mean(est_altitude(10:N10))];

clear alt15_add est_alt15_add spd15_add
clear altitude est_altitude speed

load('FlightID24234.mat')
Te = 1311/127 ; % 10.3 s
alt15_add = [altitude(1:N10)/mean(altitude(10:N10)); NaN10];
alt15_add(1:end) = alt15_add(t_index);
alt15 = [alt15 alt15_add];
est_alt15_add = [est_altitude(1:N10)/mean(est_altitude(10:N10)) ; NaN10];
est_alt15_add = est_alt15_add(t_index);
est_alt15 = [est_alt15 est_alt15_add];
spd15_add =[speed(1:N10)/mean(speed(1:N10)) ; NaN10] ;
spd15_add = spd15_add(t_index);
spd15 = [spd15 spd15_add];

residuals_factor_gps = [residuals_factor_gps mean(altitude(10:N10)) ];
residuals_factor_est = [residuals_factor_est mean(est_altitude(10:N10))];

clear alt15_add est_alt15_add spd15_add
clear altitude est_altitude speed

clear altitude speed time est_altitude;

load('FlightID30960.mat')
Te = 1123 / 106 ; % 10.6 s
alt15_add = [altitude(1:N10)/mean(altitude(10:N10)); NaN10];
alt15_add(1:end) = alt15_add(t_index);
alt15 = [alt15 alt15_add];
est_alt15_add = [est_altitude(1:N10)/mean(est_altitude(10:N10)) ; NaN10];
est_alt15_add = est_alt15_add(t_index);
est_alt15 = [est_alt15 est_alt15_add];
spd15_add =[speed(1:N10)/mean(speed(1:N10)) ; NaN10] ;
spd15_add = spd15_add(t_index);
spd15 = [spd15 spd15_add];

residuals_factor_gps = [residuals_factor_gps mean(altitude(10:N10)) ];
residuals_factor_est = [residuals_factor_est mean(est_altitude(10:N10))];

clear alt15_add est_alt15_add spd15_add
clear altitude est_altitude speed

load('FlightID31210.mat')
Te = 728/76 ;  % 9.57 s
alt15_add = [altitude(1:N10)/mean(altitude(10:N10)); NaN10];
alt15_add(1:end) = alt15_add(t_index);
alt15 = [alt15 alt15_add];
est_alt15_add = [est_altitude(1:N10)/mean(est_altitude(10:N10)) ; NaN10];
est_alt15_add = est_alt15_add(t_index);
est_alt15 = [est_alt15 est_alt15_add];
spd15_add =[speed(1:N10)/mean(speed(1:N10)) ; NaN10] ;
spd15_add = spd15_add(t_index);
spd15 = [spd15 spd15_add];

residuals_factor_gps = [residuals_factor_gps mean(altitude(10:N10)) ];
residuals_factor_est = [residuals_factor_est mean(est_altitude(10:N10))];

clear alt15_add est_alt15_add spd15_add
clear altitude est_altitude speed

load('FlightID31321.mat')
Te = 1794/179 ; %  10.0 s
alt15_add = [altitude(1:N10)/mean(altitude(10:N10)); NaN10];
alt15_add(1:end) = alt15_add(t_index);
alt15 = [alt15 alt15_add];
est_alt15_add = [est_altitude(1:N10)/mean(est_altitude(10:N10)) ; NaN10];
est_alt15_add = est_alt15_add(t_index);
est_alt15 = [est_alt15 est_alt15_add];
spd15_add =[speed(1:N10)/mean(speed(1:N10)) ; NaN10] ;
spd15_add = spd15_add(t_index);
spd15 = [spd15 spd15_add];

residuals_factor_gps = [residuals_factor_gps mean(altitude(10:N10)) ];
residuals_factor_est = [residuals_factor_est mean(est_altitude(10:N10))];

clear alt15_add est_alt15_add spd15_add
clear altitude est_altitude speed

load('FlightID31599.mat')
Te = 688/67 ; % 10.2 s
alt15_add = [altitude(1:N10)/mean(altitude(10:N10)); NaN10];
alt15_add(1:end) = alt15_add(t_index);
alt15 = [alt15 alt15_add];
est_alt15_add = [est_altitude(1:N10)/mean(est_altitude(10:N10)) ; NaN10];
est_alt15_add = est_alt15_add(t_index);
est_alt15 = [est_alt15 est_alt15_add];
spd15_add =[speed(1:N10)/mean(speed(1:N10)) ; NaN10] ;
spd15_add = spd15_add(t_index);
spd15 = [spd15 spd15_add];

residuals_factor_gps = [residuals_factor_gps mean(altitude(10:N10)) ];
residuals_factor_est = [residuals_factor_est mean(est_altitude(10:N10))];

clear alt15_add est_alt15_add spd15_add
clear altitude est_altitude speed

load('FlightID31861.mat')
Te = 570/58 ; % 9.8 s
alt15_add = [altitude(1:N10)/mean(altitude(10:N10)); NaN10];
alt15_add(1:end) = alt15_add(t_index);
alt15 = [alt15 alt15_add];
est_alt15_add = [est_altitude(1:N10)/mean(est_altitude(10:N10)) ; NaN10];
est_alt15_add = est_alt15_add(t_index);
est_alt15 = [est_alt15 est_alt15_add];
spd15_add =[speed(1:N10)/mean(speed(1:N10)) ; NaN10] ;
spd15_add = spd15_add(t_index);
spd15 = [spd15 spd15_add];

residuals_factor_gps = [residuals_factor_gps mean(altitude(10:N10)) ];
residuals_factor_est = [residuals_factor_est mean(est_altitude(10:N10))];

clear alt15_add est_alt15_add spd15_add
clear altitude est_altitude speed

load('FlightID32591.mat')
Te = 807/78 ; % 10.3 s
alt15_add = [altitude(1:N10)/mean(altitude(10:N10)); NaN10];
alt15_add(1:end) = alt15_add(t_index);
alt15 = [alt15 alt15_add];
est_alt15_add = [est_altitude(1:N10)/mean(est_altitude(10:N10)) ; NaN10];
est_alt15_add = est_alt15_add(t_index);
est_alt15 = [est_alt15 est_alt15_add];
spd15_add =[speed(1:N10)/mean(speed(1:N10)) ; NaN10] ;
spd15_add = spd15_add(t_index);
spd15 = [spd15 spd15_add];

residuals_factor_gps = [residuals_factor_gps mean(altitude(10:N10)) ];
residuals_factor_est = [residuals_factor_est mean(est_altitude(10:N10))];

clear alt15_add est_alt15_add spd15_add
clear altitude est_altitude speed

load('FlightID38745.mat')
Te = 745 /74 ; % 10.1 s
alt15_add = [altitude(1:N10)/mean(altitude(10:N10)); NaN10];
alt15_add(1:end) = alt15_add(t_index);
alt15 = [alt15 alt15_add];
est_alt15_add = [est_altitude(1:N10)/mean(est_altitude(10:N10)) ; NaN10];
est_alt15_add = est_alt15_add(t_index);
est_alt15 = [est_alt15 est_alt15_add];
spd15_add =[speed(1:N10)/mean(speed(1:N10)) ; NaN10] ;
spd15_add = spd15_add(t_index);
spd15 = [spd15 spd15_add];

residuals_factor_gps = [residuals_factor_gps mean(altitude(10:N10)) ];
residuals_factor_est = [residuals_factor_est mean(est_altitude(10:N10))];

clear alt15_add est_alt15_add spd15_add
clear altitude est_altitude speed

figure(3)
subplot(2,1,1)
shadedErrorBar(t_sort,spd15',{@nanmedian,@madX},{'-ok','markerfacecolor','k'});
set(gca,'xticklabel',[])
set(gca,'position',[0.13 0.50 0.775 0.3411627906976745])
%xlabel('Time (s)')
ylabel('Normalized groundspeed','Interpreter','latex')
axis([-1 401 0 1.5])
text(10,1.25,'(a) GPS data (flights:18 ; gulls:9)','Interpreter','latex')
box on

subplot(2,1,2)
shadedErrorBar(t_sort,alt15',{@nanmedian,@madX},{'-ok','markerfacecolor','k'});
xlabel('Time (s)','Interpreter','latex')
ylabel('Normalized altitude','Interpreter','latex')
text(10,1.25,'(b) GPS data (flights:18 ; gulls:9)','Interpreter','latex')
axis([-1 401 -0.15 1.5])
box on

%return
figure(1)
subplot(4,1,1)
shadedErrorBar(t_sort,spd15',{@nanmedian,@madX},{'-ok','markerfacecolor','k'});
%xlabel('Time (s)'
ylabel('Normalized speed','Interpreter','latex')
axis([-1 401 0 1.5])
text(10,1.25,'(a) GPS data (flights:18 ; gulls:9)','Interpreter','latex')
box on

subplot(4,1,2)
shadedErrorBar(t_sort,alt15',{@nanmedian,@madX},{'-ok','markerfacecolor','k'});
%xlabel('Time (s)')
ylabel('Normalized alt.','Interpreter','latex')
text(10,1.25,'(b) GPS data (flights:18 ; gulls:9)','Interpreter','latex')
axis([-1 401 -0.15 1.5])
box on

%% Loading data with a direct altitude control model
% Shaded nomalized graphs
% Data at Te = 15s
N15 = 27 ; % number of points
Time15 = 0:15:(N15-1)*15;
%Time = 0:10:(N10-1)*10;
Time10 = [10:30:400 20:30:400];

t = [Time15 Time10];
[t_sort,t_index] = sort(t);
NaN15 = NaN(length(Time10),1); 

load('FlightID436_alt.mat')
Te = 1080/76 ; % 14.21 s OK
alt15_direct_alt = [altitude(1:N15)/mean(altitude(7:N15)) ; NaN15];
est_alt15_direct_alt  = [est_altitude(1:N15)/mean(est_altitude(7:N15)) ; NaN15];
spd15_direct_alt = [speed(1:N15)/mean(speed(1:N15)) ; NaN15];

alt15_direct_alt = alt15_direct_alt(t_index);
est_alt15_direct_alt = est_alt15_direct_alt(t_index);
spd15_direct_alt = spd15_direct_alt(t_index);

residuals_factor_est_direct_alt = [ mean(est_altitude(7:N15))];

clear altitude est_altitude speed

load('FlightID564_alt.mat')
Te = 1755/118 ; % 14.87 s OK
alt15_add_direct_alt = [altitude(1:N15)/mean(altitude(7:N15)); NaN15];
alt15_add_direct_alt(1:end) = alt15_add_direct_alt(t_index);
alt15_direct_alt = [alt15_direct_alt alt15_add_direct_alt];
est_alt15_add_direct_alt = [est_altitude(1:N15)/mean(est_altitude(7:N15)) ; NaN15];
est_alt15_add_direct_alt = est_alt15_add_direct_alt(t_index);
est_alt15_direct_alt = [est_alt15_direct_alt est_alt15_add_direct_alt];
spd15_add_direct_alt =[speed(1:N15)/mean(speed(1:N15)) ; NaN15] ;
spd15_add_direct_alt = spd15_add_direct_alt(t_index);
spd15_direct_alt = [spd15_direct_alt spd15_add_direct_alt];

residuals_factor_est_direct_alt = [residuals_factor_est_direct_alt mean(est_altitude(7:N15))];

clear alt15_add_direct_alt est_alt15_add_direct_alt spd15_add_direct_alt
clear altitude est_altitude speed

load('FlightID571_alt.mat')
Te = 915/59; % 15.5 s OK
alt15_add_direct_alt = [altitude(1:N15)/mean(altitude(7:N15)); NaN15];
alt15_add_direct_alt(1:end) = alt15_add_direct_alt(t_index);
alt15_direct_alt = [alt15_direct_alt alt15_add_direct_alt];
est_alt15_add_direct_alt = [est_altitude(1:N15)/mean(est_altitude(7:N15)) ; NaN15];
est_alt15_add_direct_alt = est_alt15_add_direct_alt(t_index);
est_alt15_direct_alt = [est_alt15_direct_alt est_alt15_add_direct_alt];
spd15_add_direct_alt =[speed(1:N15)/mean(speed(1:N15)) ; NaN15] ;
spd15_add_direct_alt = spd15_add_direct_alt(t_index);
spd15_direct_alt = [spd15_direct_alt spd15_add_direct_alt];

residuals_factor_est_direct_alt = [residuals_factor_est_direct_alt mean(est_altitude(7:N15))];

clear alt15_add_direct_alt est_alt15_add_direct_alt spd15_add_direct_alt
clear altitude est_altitude speed

load('FlightID5917_alt.mat')
Te = 1140/77 ; % 14.80 s OK
alt15_add_direct_alt = [altitude(1:N15)/mean(altitude(7:N15)); NaN15];
alt15_add_direct_alt(1:end) = alt15_add_direct_alt(t_index);
alt15_direct_alt = [alt15_direct_alt alt15_add_direct_alt];
est_alt15_add_direct_alt = [est_altitude(1:N15)/mean(est_altitude(7:N15)) ; NaN15];
est_alt15_add_direct_alt = est_alt15_add_direct_alt(t_index);
est_alt15_direct_alt = [est_alt15_direct_alt est_alt15_add_direct_alt];
spd15_add_direct_alt =[speed(1:N15)/mean(speed(1:N15)) ; NaN15] ;
spd15_add_direct_alt = spd15_add_direct_alt(t_index);
spd15_direct_alt = [spd15_direct_alt spd15_add_direct_alt];

residuals_factor_est_direct_alt = [residuals_factor_est_direct_alt mean(est_altitude(7:N15))];

clear alt15_add_direct_alt est_alt15_add_direct_alt spd15_add_direct_alt
clear altitude est_altitude speed

load('FlightID14477_alt.mat')
Te = 1892/120 ; % 15.76 s OK
alt15_add_direct_alt = [altitude(1:N15)/mean(altitude(7:N15)); NaN15];
alt15_add_direct_alt(1:end) = alt15_add_direct_alt(t_index);
alt15_direct_alt = [alt15_direct_alt alt15_add_direct_alt];
est_alt15_add_direct_alt = [est_altitude(1:N15)/mean(est_altitude(7:N15)) ; NaN15];
est_alt15_add_direct_alt = est_alt15_add_direct_alt(t_index);
est_alt15_direct_alt = [est_alt15_direct_alt est_alt15_add_direct_alt];
spd15_add_direct_alt =[speed(1:N15)/mean(speed(1:N15)) ; NaN15] ;
spd15_add_direct_alt = spd15_add_direct_alt(t_index);
spd15_direct_alt = [spd15_direct_alt spd15_add_direct_alt];

residuals_factor_est_direct_alt = [residuals_factor_est_direct_alt mean(est_altitude(7:N15))];

clear alt15_add_direct_alt est_alt15_add_direct_alt spd15_add_direct_alt
clear altitude est_altitude speed

load('FlightID16542_alt.mat')
Te = 1327/83 ; % 15.98 s OK
alt15_add_direct_alt = [altitude(1:N15)/mean(altitude(7:N15)); NaN15];
alt15_add_direct_alt(1:end) = alt15_add_direct_alt(t_index);
alt15_direct_alt = [alt15_direct_alt alt15_add_direct_alt];
est_alt15_add_direct_alt = [est_altitude(1:N15)/mean(est_altitude(7:N15)) ; NaN15];
est_alt15_add_direct_alt = est_alt15_add_direct_alt(t_index);
est_alt15_direct_alt = [est_alt15_direct_alt est_alt15_add_direct_alt];
spd15_add_direct_alt =[speed(1:N15)/mean(speed(1:N15)) ; NaN15] ;
spd15_add_direct_alt = spd15_add_direct_alt(t_index);
spd15_direct_alt = [spd15_direct_alt spd15_add_direct_alt];

residuals_factor_est_direct_alt = [residuals_factor_est_direct_alt mean(est_altitude(7:N15))];

clear alt15_add_direct_alt est_alt15_add_direct_alt spd15_add_direct_alt
clear altitude est_altitude speed

% Data at Te = 10s
% N = 41 ; % number of points
% alt = [];
% est_alt10 = [];
% spd = [];
clear t t_sort t_index Time15 Time10 NaN15

N10 = 41;
Time15 = 15:30:400;
Time10 = 0:10:(N10-1)*10;

t = [Time10 Time15];
[t_sort,t_index] = sort(t);
NaN10 = NaN(length(Time15),1); 

load('FlightID5225_alt.mat')
Te = 810/82 ; % 9.87s
alt15_add_direct_alt = [altitude(1:N10)/mean(altitude(10:N10)); NaN10];
alt15_add_direct_alt(1:end) = alt15_add_direct_alt(t_index);
alt15_direct_alt = [alt15_direct_alt alt15_add_direct_alt];
est_alt15_add_direct_alt = [est_altitude(1:N10)/mean(est_altitude(10:N10)) ; NaN10];
est_alt15_add_direct_alt = est_alt15_add_direct_alt(t_index);
est_alt15_direct_alt = [est_alt15_direct_alt est_alt15_add_direct_alt];
spd15_add_direct_alt =[speed(1:N10)/mean(speed(1:N10)) ; NaN10] ;
spd15_add_direct_alt = spd15_add_direct_alt(t_index);
spd15_direct_alt = [spd15_direct_alt spd15_add_direct_alt];

residuals_factor_est_direct_alt = [residuals_factor_est_direct_alt mean(est_altitude(10:N10))];

clear alt15_add_direct_alt est_alt15_add_direct_alt spd15_add_direct_alt
clear altitude est_altitude speed

load('FlightID24611_alt.mat')
Te = 10.29 ; 
alt15_add_direct_alt = [altitude(1:N10)/mean(altitude(10:N10)); NaN10];
alt15_add_direct_alt(1:end) = alt15_add_direct_alt(t_index);
alt15_direct_alt = [alt15_direct_alt alt15_add_direct_alt];
est_alt15_add_direct_alt = [est_altitude(1:N10)/mean(est_altitude(10:N10)) ; NaN10];
est_alt15_add_direct_alt = est_alt15_add_direct_alt(t_index);
est_alt15_direct_alt = [est_alt15_direct_alt est_alt15_add_direct_alt];
spd15_add_direct_alt =[speed(1:N10)/mean(speed(1:N10)) ; NaN10] ;
spd15_add_direct_alt = spd15_add_direct_alt(t_index);
spd15_direct_alt = [spd15_direct_alt spd15_add_direct_alt];

residuals_factor_est_direct_alt = [residuals_factor_est_direct_alt mean(est_altitude(10:N10))];

clear alt15_add_direct_alt est_alt15_add_direct_alt spd15_add_direct_alt
clear altitude est_altitude speed

load('FlightID16606_alt.mat')
Te = 887/85 ; % 10.4 s
alt15_add_direct_alt = [altitude(1:N10)/mean(altitude(10:N10)); NaN10];
alt15_add_direct_alt(1:end) = alt15_add_direct_alt(t_index);
alt15_direct_alt = [alt15_direct_alt alt15_add_direct_alt];
est_alt15_add_direct_alt = [est_altitude(1:N10)/mean(est_altitude(10:N10)) ; NaN10];
est_alt15_add_direct_alt = est_alt15_add_direct_alt(t_index);
est_alt15_direct_alt = [est_alt15_direct_alt est_alt15_add_direct_alt];
spd15_add_direct_alt =[speed(1:N10)/mean(speed(1:N10)) ; NaN10] ;
spd15_add_direct_alt = spd15_add_direct_alt(t_index);
spd15_direct_alt = [spd15_direct_alt spd15_add_direct_alt];

residuals_factor_est_direct_alt = [residuals_factor_est_direct_alt mean(est_altitude(10:N10))];

clear alt15_add_direct_alt est_alt15_add_direct_alt spd15_add_direct_alt
clear altitude est_altitude speed

load('FlightID20104_alt.mat')
Te = 2165 /208 ; % 10.4 s
alt15_add_direct_alt = [altitude(1:N10)/mean(altitude(10:N10)); NaN10];
alt15_add_direct_alt(1:end) = alt15_add_direct_alt(t_index);
alt15_direct_alt = [alt15_direct_alt alt15_add_direct_alt];
est_alt15_add_direct_alt = [est_altitude(1:N10)/mean(est_altitude(10:N10)) ; NaN10];
est_alt15_add_direct_alt = est_alt15_add_direct_alt(t_index);
est_alt15_direct_alt = [est_alt15_direct_alt est_alt15_add_direct_alt];
spd15_add_direct_alt =[speed(1:N10)/mean(speed(1:N10)) ; NaN10] ;
spd15_add_direct_alt = spd15_add_direct_alt(t_index);
spd15_direct_alt = [spd15_direct_alt spd15_add_direct_alt];

residuals_factor_est_direct_alt = [residuals_factor_est_direct_alt mean(est_altitude(10:N10))];

clear alt15_add_direct_alt est_alt15_add_direct_alt spd15_add_direct_alt
clear altitude est_altitude speed

load('FlightID24234_alt.mat')
Te = 1311/127 ; % 10.3 s
alt15_add_direct_alt = [altitude(1:N10)/mean(altitude(10:N10)); NaN10];
alt15_add_direct_alt(1:end) = alt15_add_direct_alt(t_index);
alt15_direct_alt = [alt15_direct_alt alt15_add_direct_alt];
est_alt15_add_direct_alt = [est_altitude(1:N10)/mean(est_altitude(10:N10)) ; NaN10];
est_alt15_add_direct_alt = est_alt15_add_direct_alt(t_index);
est_alt15_direct_alt = [est_alt15_direct_alt est_alt15_add_direct_alt];
spd15_add_direct_alt =[speed(1:N10)/mean(speed(1:N10)) ; NaN10] ;
spd15_add_direct_alt = spd15_add_direct_alt(t_index);
spd15_direct_alt = [spd15_direct_alt spd15_add_direct_alt];

residuals_factor_est_direct_alt = [residuals_factor_est_direct_alt mean(est_altitude(10:N10))];

clear alt15_add_direct_alt est_alt15_add_direct_alt spd15_add_direct_alt
clear altitude est_altitude speed

load('FlightID30960_alt.mat')
Te = 1123 / 106 ; % 10.6 s
alt15_add_direct_alt = [altitude(1:N10)/mean(altitude(10:N10)); NaN10];
alt15_add_direct_alt(1:end) = alt15_add_direct_alt(t_index);
alt15_direct_alt = [alt15_direct_alt alt15_add_direct_alt];
est_alt15_add_direct_alt = [est_altitude(1:N10)/mean(est_altitude(10:N10)) ; NaN10];
est_alt15_add_direct_alt = est_alt15_add_direct_alt(t_index);
est_alt15_direct_alt = [est_alt15_direct_alt est_alt15_add_direct_alt];
spd15_add_direct_alt =[speed(1:N10)/mean(speed(1:N10)) ; NaN10] ;
spd15_add_direct_alt = spd15_add_direct_alt(t_index);
spd15_direct_alt = [spd15_direct_alt spd15_add_direct_alt];

residuals_factor_est_direct_alt = [residuals_factor_est_direct_alt mean(est_altitude(10:N10))];

clear alt15_add_direct_alt est_alt15_add_direct_alt spd15_add_direct_alt
clear altitude est_altitude speed

load('FlightID31210_alt.mat')
Te = 728/76 ;  % 9.57 s
alt15_add_direct_alt = [altitude(1:N10)/mean(altitude(10:N10)); NaN10];
alt15_add_direct_alt(1:end) = alt15_add_direct_alt(t_index);
alt15_direct_alt = [alt15_direct_alt alt15_add_direct_alt];
est_alt15_add_direct_alt = [est_altitude(1:N10)/mean(est_altitude(10:N10)) ; NaN10];
est_alt15_add_direct_alt = est_alt15_add_direct_alt(t_index);
est_alt15_direct_alt = [est_alt15_direct_alt est_alt15_add_direct_alt];
spd15_add_direct_alt =[speed(1:N10)/mean(speed(1:N10)) ; NaN10] ;
spd15_add_direct_alt = spd15_add_direct_alt(t_index);
spd15_direct_alt = [spd15_direct_alt spd15_add_direct_alt];

residuals_factor_est_direct_alt = [residuals_factor_est_direct_alt mean(est_altitude(10:N10))];

clear alt15_add_direct_alt est_alt15_add_direct_alt spd15_add_direct_alt
clear altitude est_altitude speed

load('FlightID31321_alt.mat')
Te = 1794/179 ; %  10.0 s
alt15_add_direct_alt = [altitude(1:N10)/mean(altitude(10:N10)); NaN10];
alt15_add_direct_alt(1:end) = alt15_add_direct_alt(t_index);
alt15_direct_alt = [alt15_direct_alt alt15_add_direct_alt];
est_alt15_add_direct_alt = [est_altitude(1:N10)/mean(est_altitude(10:N10)) ; NaN10];
est_alt15_add_direct_alt = est_alt15_add_direct_alt(t_index);
est_alt15_direct_alt = [est_alt15_direct_alt est_alt15_add_direct_alt];
spd15_add_direct_alt =[speed(1:N10)/mean(speed(1:N10)) ; NaN10] ;
spd15_add_direct_alt = spd15_add_direct_alt(t_index);
spd15_direct_alt = [spd15_direct_alt spd15_add_direct_alt];

residuals_factor_est_direct_alt = [residuals_factor_est_direct_alt mean(est_altitude(10:N10))];

clear alt15_add_direct_alt est_alt15_add_direct_alt spd15_add_direct_alt
clear altitude est_altitude speed

load('FlightID31599_alt.mat')
Te = 688/67 ; % 10.2 s
alt15_add_direct_alt = [altitude(1:N10)/mean(altitude(10:N10)); NaN10];
alt15_add_direct_alt(1:end) = alt15_add_direct_alt(t_index);
alt15_direct_alt = [alt15_direct_alt alt15_add_direct_alt];
est_alt15_add_direct_alt = [est_altitude(1:N10)/mean(est_altitude(10:N10)) ; NaN10];
est_alt15_add_direct_alt = est_alt15_add_direct_alt(t_index);
est_alt15_direct_alt = [est_alt15_direct_alt est_alt15_add_direct_alt];
spd15_add_direct_alt =[speed(1:N10)/mean(speed(1:N10)) ; NaN10] ;
spd15_add_direct_alt = spd15_add_direct_alt(t_index);
spd15_direct_alt = [spd15_direct_alt spd15_add_direct_alt];

residuals_factor_est_direct_alt = [residuals_factor_est_direct_alt mean(est_altitude(10:N10))];

clear alt15_add_direct_alt est_alt15_add_direct_alt spd15_add_direct_alt
clear altitude est_altitude speed

load('FlightID31861_alt.mat')
Te = 570/58 ; % 9.8 s
alt15_add_direct_alt = [altitude(1:N10)/mean(altitude(10:N10)); NaN10];
alt15_add_direct_alt(1:end) = alt15_add_direct_alt(t_index);
alt15_direct_alt = [alt15_direct_alt alt15_add_direct_alt];
est_alt15_add_direct_alt = [est_altitude(1:N10)/mean(est_altitude(10:N10)) ; NaN10];
est_alt15_add_direct_alt = est_alt15_add_direct_alt(t_index);
est_alt15_direct_alt = [est_alt15_direct_alt est_alt15_add_direct_alt];
spd15_add_direct_alt =[speed(1:N10)/mean(speed(1:N10)) ; NaN10] ;
spd15_add_direct_alt = spd15_add_direct_alt(t_index);
spd15_direct_alt = [spd15_direct_alt spd15_add_direct_alt];

residuals_factor_est_direct_alt = [residuals_factor_est_direct_alt mean(est_altitude(10:N10))];

clear alt15_add_direct_alt est_alt15_add_direct_alt spd15_add_direct_alt
clear altitude est_altitude speed

load('FlightID32591_alt.mat')
Te = 807/78 ; % 10.3 s
alt15_add_direct_alt = [altitude(1:N10)/mean(altitude(10:N10)); NaN10];
alt15_add_direct_alt(1:end) = alt15_add_direct_alt(t_index);
alt15_direct_alt = [alt15_direct_alt alt15_add_direct_alt];
est_alt15_add_direct_alt = [est_altitude(1:N10)/mean(est_altitude(10:N10)) ; NaN10];
est_alt15_add_direct_alt = est_alt15_add_direct_alt(t_index);
est_alt15_direct_alt = [est_alt15_direct_alt est_alt15_add_direct_alt];
spd15_add_direct_alt =[speed(1:N10)/mean(speed(1:N10)) ; NaN10] ;
spd15_add_direct_alt = spd15_add_direct_alt(t_index);
spd15_direct_alt = [spd15_direct_alt spd15_add_direct_alt];

residuals_factor_est_direct_alt = [residuals_factor_est_direct_alt mean(est_altitude(10:N10))];

clear alt15_add_direct_alt est_alt15_add_direct_alt spd15_add_direct_alt
clear altitude est_altitude speed

load('FlightID38745_alt.mat')
Te = 745 /74 ; % 10.1 s
alt15_add_direct_alt = [altitude(1:N10)/mean(altitude(10:N10)); NaN10];
alt15_add_direct_alt(1:end) = alt15_add_direct_alt(t_index);
alt15_direct_alt = [alt15_direct_alt alt15_add_direct_alt];
est_alt15_add_direct_alt = [est_altitude(1:N10)/mean(est_altitude(10:N10)) ; NaN10];
est_alt15_add_direct_alt = est_alt15_add_direct_alt(t_index);
est_alt15_direct_alt = [est_alt15_direct_alt est_alt15_add_direct_alt];
spd15_add_direct_alt =[speed(1:N10)/mean(speed(1:N10)) ; NaN10] ;
spd15_add_direct_alt = spd15_add_direct_alt(t_index);
spd15_direct_alt = [spd15_direct_alt spd15_add_direct_alt];

residuals_factor_est_direct_alt = [residuals_factor_est_direct_alt mean(est_altitude(10:N10))];

clear alt15_add_direct_alt est_alt15_add_direct_alt spd15_add_direct_alt
clear altitude est_altitude speed

%%
subplot(4,1,3)
shadedErrorBar(t_sort,est_alt15_direct_alt',{@nanmedian,@madX},{'-or','markerfacecolor','r'});
%xlabel('Time (s)')
ylabel('Normalized alt.','Interpreter','latex')
text(10,1.25,'(c) Altitude control model','Interpreter','latex')
axis([-1 401 -0.15 1.5])
box on


%% Graph optic flow-based control model
subplot(4,1,4)
shadedErrorBar(t_sort,est_alt15',{@nanmedian,@madX},{'-ob','markerfacecolor','b'});
xlabel('Time (s)','Interpreter','latex')
ylabel('Normalized alt.','Interpreter','latex')
text(10,1.25,'(d) Optic flow-based control model','Interpreter','latex')
axis([-1 401 -0.15 1.5])
box on

%print -f1 -dpng -r600 Fig_all_gulls
%print -f3 -dpng -r600 Fig_all_gulls_GPSdata

return

%%%%%%%%%%%%%%% FIN %%%%%%%%%%%%%
% figure(2)
% subplot(2,1,1)
% x = Time;
% y = mean(spd');
% err = std(spd');
% errorbar(x,y,err,'-o','MarkerSize',5,'MarkerEdgeColor',[0 0.4470 0.7410],'MarkerFaceColor',[0 0.4470 0.7410])
% %xlabel('Time (s)')
% ylabel('Normalized speed')
% axis([-1 401 0.3 1.3])
% box on
% 
% subplot(2,1,2)
% x = Time;
% y = mean(alt');
% err = std(alt');
% errorbar(x,y,err,'-o','MarkerSize',5,'MarkerEdgeColor',[0 0.447 0.741],'MarkerFaceColor',[0 0.4470 0.7410])
% xlabel('Time (s)')
% ylabel('Normalized altitude')
% axis([-1 401 0 1.5])
% box on
% 
% %print('fig_multi_graph','-f2','-dpng','-r600')
% 
% % Superpostion of the 2 sets of data Te = 15 s & Te = 10 s
% 
% %Time15(1:2:end) % list every 30s
% %Time(1:3:end)
% 
% x = Time;
% % adding commons point inside the mean
% y = mean(alt');
% y15 = mean(alt15');
% 
% y(1:3:end) = (y(1:3:end)*12+y15(1:2:end)*6)/18;
% % adding commons point inside the sdt (corrected sd)
% err = std(alt');
% y15 = std(alt15');
% err(1:3:end) = sqrt(((err(1:3:end).*err(1:3:end))*11+y15(1:2:end).*y15(1:2:end)*5)/17);
% hold on
% errorbar(x,y,err,'-o','MarkerSize',5,'MarkerEdgeColor','r','MarkerFaceColor','r','Color','r')
% hold on
% y15mean = mean(alt15');
% y15std = std(alt15');
% errorbar(Time15(2:2:end),y15mean(2:2:end),y15std(2:2:end),'o','MarkerSize',5,'MarkerEdgeColor','m','MarkerFaceColor','m','Color','m')
% 
% % including the 2 sets of data
% figure(3)
% t = [Time Time15(2:2:end)];
% [t_sort,t_index] = sort(t);
% 
% subplot(3,1,1)
% v = mean(spd');
% v15 = mean(spd15');
% v(1:3:end) = (v(1:3:end)*12+v15(1:2:end)*6)/18;
% 
% v_err = std(spd');
% v15_err = std(spd15');
% v_err(1:3:end) = sqrt(((err(1:3:end).*err(1:3:end))*11+v15_err(1:2:end).*v15_err(1:2:end)*5)/17);
% 
% vmean_sort = [v v15(2:2:end)];
% vmean_sort(1:end) = vmean_sort(t_index);
% vstd_sort = [v_err v15_err(2:2:end)];
% vstd_sort(1:end) = vstd_sort(t_index); 
% errorbar(t_sort,vmean_sort,vstd_sort,'-o','MarkerSize',5,'MarkerEdgeColor',[0 0.447 0.741],'MarkerFaceColor',[0 0.4470 0.7410])
% ylabel('Normalized speed')
% axis([-1 401 0.3 1.3])
% box on
% 
% subplot(3,1,2)
% ymean_sort = [y y15mean(2:2:end)];
% ymean_sort(1:end) = ymean_sort(t_index);
% ystd_sort = [err y15std(2:2:end)];
% ystd_sort(1:end) = ystd_sort(t_index); 
% errorbar(t_sort,ymean_sort,ystd_sort,'-o','MarkerSize',5,'MarkerEdgeColor',[0 0.447 0.741],'MarkerFaceColor',[0 0.4470 0.7410])
% xlabel('Time (s)')
% ylabel('Normalized altitude')
% axis([-1 401 0 1.5])
% box on
% 
% subplot(3,1,3)
% est_z = mean(est_alt10');
% est_z15 = mean(est_alt15');
% est_z(1:3:end) = (est_z(1:3:end)*12+est_z15(1:2:end)*6)/18;
% 
% z_err = std(est_alt10');
% z15_err = std(est_alt15');
% z_err(1:3:end) = sqrt(((z_err(1:3:end).*z_err(1:3:end))*11+z15_err(1:2:end).*z15_err(1:2:end)*5)/17);
% 
% zmean_sort = [est_z est_z15(2:2:end)];
% zmean_sort(1:end) = zmean_sort(t_index);
% zstd_sort = [z_err z15_err(2:2:end)];
% zstd_sort(1:end) = zstd_sort(t_index); 
% errorbar(t_sort,zmean_sort,zstd_sort,'-o','MarkerSize',5,'MarkerEdgeColor',[0 0.447 0.741],'MarkerFaceColor',[0 0.4470 0.7410])
% ylabel('Normalized estimated alt.')
% axis([-1 401 0 1.5])
% box on
% 
% return
% 
% figure(2)
% subplot(2,1,1)
% x = Time;
% y = mean(spd');
% err = std(spd');
% errorbar(x,y,err,'-o','MarkerSize',5,'MarkerEdgeColor',[0 0.4470 0.7410],'MarkerFaceColor',[0 0.4470 0.7410])
% %xlabel('Time (s)')
% ylabel('Normalized speed')
% axis([-1 401 0.3 1.3])
% box on
% 
% subplot(2,1,2)
% x = Time;
% y = mean(alt');
% err = std(alt');
% errorbar(x,y,err,'-o','MarkerSize',5,'MarkerEdgeColor',[0 0.447 0.741],'MarkerFaceColor',[0 0.4470 0.7410])
% xlabel('Time (s)')
% ylabel('Normalized altitude')
% axis([-1 401 0 1.5])
% box on
```
