## Supplementary material for "Optic flow helps explain gulls’ altitude control over seas": data set of gulls takeoffs - Flight modelling: multi_graph3.m

```
% Figure tra�ant les erreurs des mod�les
clear all
clc
close all
multi_graph2

%

figure(2)
subplot(2,1,1)
shadedErrorBar(t_sort,(alt15-est_alt15_direct_alt)',{@nanmedian,@madX},{'-or','markerfacecolor', [0.1 0.5 0.7]});
%xlabel('Time (s)')
ylabel('Modelling error','Interpreter','latex')
axis([-1 401 -0.4 0.4])
text(10,0.3,'(a) between GPS data and altitude control model (flights:18 ; gulls:9)','Interpreter','latex')
box on

subplot(2,1,2)
shadedErrorBar(t_sort,(alt15-est_alt15)',{@nanmedian,@madX},{'-ob','markerfacecolor','b'});
xlabel('Time (s)','Interpreter','latex')
ylabel('Modelling error','Interpreter','latex')
text(10,0.3,'(b) between GPS data and optic flow-based control model (flights:18 ; gulls:9)','Interpreter','latex')
axis([-1 401 -0.4 0.4])
box on


figure(4)
subplot(4,1,1)
h = area([200 200 401 401],[-0.15 1.5 1.5 -0.15])
set(h,'FaceColor',[0.5 0.5 0.5])
set(h,'EdgeColor','none')
set(h,'EdgeAlpha',0.1)
set(h,'FaceAlpha',0.1)
set(h,'ShowBaseLine','off')
hold on
h = area([200 200 401 401],[0 -0.15 -0.15 0])
set(h,'FaceColor',[0.5 0.5 0.5])
set(h,'EdgeColor','none')
set(h,'EdgeAlpha',0.1)
set(h,'FaceAlpha',0.1)
set(h,'ShowBaseLine','off')
hold on
%patch([201 201 400 400],[-0.1 1.45 1.45 -0.1],[0.9 0.9 0.9],'FaceAlpha',0.5,'EdgeColor',[0.9 0.9 0.9])

shadedErrorBar(t_sort(1:29),est_alt15_direct_alt(1:29,:)',{@nanmedian,@madX},{'-or','markerfacecolor','r'});
hold on
shadedErrorBar(t_sort(29:end),est_alt15_direct_alt(29:end,:)',{@nanmedian,@madX},{'-or','markerfacecolor',[0.7 0 0]});
set(gca,'xticklabel',[])
%xlabel('Time (s)')
ylabel('Normalized alt.','FontSize', 9,'Interpreter','latex')
text(295, 1.7, '\textbf{flights:18 ; gulls:9}','Interpreter','latex')
text(0,1.7,'(a) Altitude control model','Interpreter','latex')
axis([-1 401 -0.15 1.5])
box on

subplot(4,1,2)
h = area([200 200 401 401],[-0.15 1.5 1.5 -0.15])
set(h,'FaceColor',[0.5 0.5 0.5])
set(h,'EdgeColor','none')
set(h,'EdgeAlpha',0.1)
set(h,'FaceAlpha',0.1)
set(h,'ShowBaseLine','off')
hold on
h = area([200 200 401 401],[0 -0.15 -0.15 0])
set(h,'FaceColor',[0.5 0.5 0.5])
set(h,'EdgeColor','none')
set(h,'EdgeAlpha',0.1)
set(h,'FaceAlpha',0.1)
set(h,'ShowBaseLine','off')
hold on
shadedErrorBar(t_sort(1:29),est_alt15(1:29,:)',{@nanmedian,@madX},{'-ob','markerfacecolor','b'});
hold on
shadedErrorBar(t_sort(29:end),est_alt15(29:end,:)',{@nanmedian,@madX},{'-ob','markerfacecolor',[0 0 0.7]});
set(gca,'xticklabel',[])
%xlabel('Time (s)','Interpreter','latex')
ylabel('Normalized alt.','FontSize', 9,'Interpreter','latex')
text(0,1.7,'(b) Optic flow-based control model','Interpreter','latex')
axis([-1 401 -0.15 1.5])
box on


subplot(4,1,3)
%h = area([200 200 401 401],[-0.41 1.5 1.5 -0.41])
h = area([200 200 401 401],[-16 16 16 -16])
set(h,'FaceColor',[0.5 0.5 0.5])
set(h,'EdgeColor','none')
set(h,'EdgeAlpha',0.1)
set(h,'FaceAlpha',0.1)
set(h,'ShowBaseLine','off')
hold on
%h = area([200 200 401 401],[0 -0.41 -0.41 0])
h = area([200 200 401 401],[0 -16 -16 0])
set(h,'FaceColor',[0.5 0.5 0.5])
set(h,'EdgeColor','none')
set(h,'EdgeAlpha',0.1)
set(h,'FaceAlpha',0.1)
set(h,'ShowBaseLine','off')
hold on
shadedErrorBar(t_sort(1:29),(alt15(1:29,:).*residuals_factor_gps - est_alt15_direct_alt(1:29,:).*residuals_factor_est_direct_alt)',{@nanmedian,@madX},{'-or','markerfacecolor','r'});
hold on
shadedErrorBar(t_sort(29:end),(alt15(29:end,:).*residuals_factor_gps - est_alt15_direct_alt(29:end,:).*residuals_factor_est_direct_alt)',{@nanmedian,@madX},{'-or','markerfacecolor',[0.7 0 0]});
set(gca,'xticklabel',[])
%xlabel('Time (s)')
%xlabel('Time (s)','Interpreter','latex')
text(0,21,'(c) between GPS data and altitude control model (median $\pm$ mad)','Interpreter','latex')
ylabel('Residuals (m)','FontSize', 9,'Interpreter','latex')
text(75,10,'-4.420 $\pm$ 1.325','Interpreter','latex')
text(275,10,'-0.765 $\pm$ 0.534','Interpreter','latex')

axis([-1 401 -16 16])
box on

% shadedErrorBar(t_sort(1:29),(alt15(1:29,:)-est_alt15_direct_alt(1:29,:))',{@nanmedian,@madX},{'-or','markerfacecolor','r'});
% hold on
% shadedErrorBar(t_sort(29:end),(alt15(29:end,:)-est_alt15_direct_alt(29:end,:))',{@nanmedian,@madX},{'-or','markerfacecolor',[0.7 0 0]});
% set(gca,'xticklabel',[])
% %xlabel('Time (s)')
% ylabel('Residuals','Interpreter','latex')
% axis([-1 401 -0.41 0.41])
% text(10,0.5,'(c) between GPS data and altitude control model','Interpreter','latex')
% box on

subplot(4,1,4)
h = area([200 200 401 401],[-16 16 16 -16])
set(h,'FaceColor',[0.5 0.5 0.5])
set(h,'EdgeColor','none')
set(h,'EdgeAlpha',0.1)
set(h,'FaceAlpha',0.1)
set(h,'ShowBaseLine','off')
hold on
h = area([200 200 401 401],[0 -16 -16 0])
set(h,'FaceColor',[0.5 0.5 0.5])
set(h,'EdgeColor','none')
set(h,'EdgeAlpha',0.1)
set(h,'FaceAlpha',0.1)
set(h,'ShowBaseLine','off')
hold on
shadedErrorBar(t_sort(1:29),(alt15(1:29,:).*residuals_factor_gps - est_alt15(1:29,:).*residuals_factor_est)',{@nanmedian,@madX},{'-ob','markerfacecolor','b'});
hold on
shadedErrorBar(t_sort(29:end),(alt15(29:end,:).*residuals_factor_gps - est_alt15(29:end,:).*residuals_factor_est)',{@nanmedian,@madX},{'-ob','markerfacecolor',[0 0 0.7]});
xlabel('Time (s)','Interpreter','latex')
text(0,21,'(d) between GPS data and optic flow-based control model (median $\pm$ mad)','Interpreter','latex')
ylabel('Residuals (m)','FontSize', 9,'Interpreter','latex')
text(75,10,'0.012 $\pm$ 1.123','Interpreter','latex')
text(275,10,'0.584 $\pm$ 0.655','Interpreter','latex')
axis([-1 401 -16 16])
box on

% shadedErrorBar(t_sort(1:29),(alt15(1:29,:)-est_alt15(1:29,:))',{@nanmedian,@madX},{'-ob','markerfacecolor','b'});
% hold on
% shadedErrorBar(t_sort(29:end),(alt15(29:end,:)-est_alt15(29:end,:))',{@nanmedian,@madX},{'-ob','markerfacecolor',[0 0 0.7]});
% xlabel('Time (s)','Interpreter','latex')
% ylabel('Residuals','Interpreter','latex')
% text(10,0.5,'(d) between GPS data and optic flow-based control model','Interpreter','latex')
% axis([-1 401 -0.41 0.41])
% box on

print -f2 -dpng -r600 Fig_all_gulls_errors_models
print -f4 -dpng -r600 Fig_errors_and_models

% Stats on modelling errors
%(alt15-est_alt15)'
```
