## Supplementary material for "Optic flow helps explain gulls’ altitude control over seas": data set of gulls takeoffs - Flight modelling: multi_graph4.m

```
% Figure traçant les histogrammes des erreurs

figure(5)
% map = brewermap(2,'Set1'); 
% histf(nanmedian((alt15-est_alt15)'),-.15:.015:0.15,'facecolor',map(1,:),'facealpha',.5,'edgecolor','none')
% hold on
% histf(nanmedian((alt15-est_alt15_direct_alt)'),-.15:.015:0.15,'facecolor',map(2,:),'facealpha',.5,'edgecolor','none')
% box off
% axis tight
% %legalpha('H1','H2','location','northwest')
% legend boxoff
x1 = nanmedian((alt15.*residuals_factor_gps-est_alt15.*residuals_factor_est)')';
x2 = nanmedian((alt15.*residuals_factor_gps-est_alt15_direct_alt.*residuals_factor_est_direct_alt)')';


boxplot([x1(1:27),x2(1:27),x1(28:54),x2(28:54)],'Labels',{'OF-based','Altitude-based','OF-based','Altitude-based'},'Whisker',2)
ylabel('Residuals (m)','FontSize',12)

h = findobj(gca,'Tag','Box');

 for j=1:2
    patch(get(h(j),'XData'),get(h(j),'YData'),'y','FaceColor',[0.7 0.7 0.7]);
 end
hold on 
boxplot([x1(1:27),x2(1:27),x1(28:54),x2(28:54)],'Labels',{'OF-based','Altitude-based','OF-based','Altitude-based'},'Whisker',2)


yt = get(gca, 'YTick');
xt = get(gca, 'XTick');
hold on
plot(xt([1 2]), [1 1]*max(yt)*1.8, '-k')
text(1.4, max(yt)*1.9, '***','FontSize',16)
text(0.9, 5, 'Transient response','FontSize',12)

hold on
plot(xt([3 4]), [1 1]*max(yt)*1.8, '-k')
text(3.4, max(yt)*1.9,'***','FontSize',16)
text(2.8, 5,'Steady-state response','FontSize',12)
axis([0.5 4.5 -9 6])

print -f5 -dpng -r600 Residuals_boxplot

[p,h] = ranksum(x1(1:27),x2(1:27))
[p,h] = lillietest(x1(1:27)) % p>0.05
[p,h] = lillietest(x2(1:27)) % p>0.05
[p,h] = lillietest(x1(28:54)) % p>0.05
[p,h] = lillietest(x2(28:54)) % p<0.05 un jeu non normal
% Test non paramétrique
[p,h] = ranksum(x1(1:27),x2(1:27),'tail','right')
[p,h] = ranksum(x1(28:54),x2(28:54),'tail','right')

[h,p] = ttest(x1(1:27),x2(1:27),'tail','right')
[h,p] = ttest(x1(28:54),x2(28:54),'tail','left')

[h,p] = ttest(x1(28:54),0)
[h,p] = ttest(x1(1:27),0)
% Dans les 2 cas la distribution est centrée autour de 0

[h,p] = ttest(x2(28:54),0)
[h,p] = ttest(x2(1:27),0)
% Dans les 2 cas la distribution n'est pas centrée autour de 0

[h,p] = ranksum(x1(1:27),x2(1:27),'tail','right')
% donc gauche qui est plus élevé
[h,p] = ranksum(x1(28:54),x2(28:54),'tail','right')
% donc gauche qui est plus élevé

%[p,h] = ranksum(x,y)
%[h,p] = ttest(___)
[h,p] = ranksum(x1(1:27)-x2(1:27),2.77,'tail','right')
% p = 0.19, non significatif
% 2.77 > mean(x1 - x2)
[h,p] = ttest(x1(1:27)-x2(1:27),2.77,'tail','right')
% Test the null hypothesis that the data comes 
%  the null hypothesis at the 5% significance level,
% in favor of the alternate hypothesis that the data comes
% from a distribution with a mean greater than 2.77. 
[h,p] = ttest(x1(28:54)-x2(28:54),2.77,'tail','right')

[rho1,pvalue1] = corrcoef(x1(1:27), x2(1:27))
[rho2,pvalue2] = corrcoef(x1(28:54), x2(28:54))
```
