## Supplementary material for "Optic flow helps explain gulls’ altitude control over seas": data set of gulls takeoffs - Flight modelling: multi_graph5.m

```
% Figure traçant les histogrammes des résidus en mètres
warning off
clc
figure(6)

%alt15 = alt15.*residuals_factor_gps;
%est_alt15 = est_alt15.*residuals_factor_est;

% Direct altitude control mean +/- sd
x0_mean = mean(nanmean((alt15.*residuals_factor_gps - est_alt15_direct_alt.*residuals_factor_est_direct_alt)')');
x0_sd =  std(nanmean((alt15.*residuals_factor_gps - est_alt15_direct_alt.*residuals_factor_est_direct_alt)')');

% Direct altitude control median +/- mad
x0to29_median = nanmedian(nanmean((alt15(1:29,:).*residuals_factor_gps - est_alt15_direct_alt(1:29,:).*residuals_factor_est_direct_alt)')');
x0to29_mad =  madX(nanmean((alt15(1:29,:).*residuals_factor_gps - est_alt15_direct_alt(1:29,:).*residuals_factor_est_direct_alt)')');

x0from29_median = nanmedian(nanmean((alt15(29:end,:).*residuals_factor_gps - est_alt15_direct_alt(29:end,:).*residuals_factor_est_direct_alt)')');
x0from29_mad =  madX(nanmean((alt15(29:end,:).*residuals_factor_gps - est_alt15_direct_alt(29:end,:).*residuals_factor_est_direct_alt)')');

% Optic flow-based control mean +/- sd
x1_mean = mean(nanmean((alt15.*residuals_factor_gps - est_alt15.*residuals_factor_est)')');
x1_std = std(nanstd((alt15.*residuals_factor_gps - est_alt15.*residuals_factor_est)')');

x1to29_median = nanmedian(nanmedian((alt15(1:29,:).*residuals_factor_gps - est_alt15(1:29,:).*residuals_factor_est)')');
x1to29_mad =  madX(nanmedian((alt15(1:29,:).*residuals_factor_gps - est_alt15(1:29,:).*residuals_factor_est)')');

x1from29_median = nanmedian(nanmedian((alt15(29:end,:).*residuals_factor_gps - est_alt15(29:end,:).*residuals_factor_est)')');
x1from29_mad =  madX(nanmedian((alt15(29:end,:).*residuals_factor_gps - est_alt15(29:end,:).*residuals_factor_est)')');


subplot(2,1,1)
shadedErrorBar(t_sort,(alt15.*residuals_factor_gps-est_alt15_direct_alt.*residuals_factor_est_direct_alt)',{@nanmean,@nanstd},{'-or','markerfacecolor',[0.1 0.5 0.7]});
%xlabel('Time (s)')
%xlabel('Time (s)','Interpreter','latex')
text(10,23,'(a) between GPS data and altitude control model (flights:18 ; gulls:9)','Interpreter','latex')
ylabel('Residuals (m)','Interpreter','latex')
%text(10,15,'(a)','Interpreter','latex')
text(45,13,'-2.626 $\pm$ 2.489 m (mean $\pm$ sd)','Interpreter','latex')
axis([-1 401 -20 20])
box on

subplot(2,1,2)
shadedErrorBar(t_sort,(alt15.*residuals_factor_gps-est_alt15.*residuals_factor_est)',{@nanmean,@nanstd},{'-ob','markerfacecolor','b'});
xlabel('Time (s)','Interpreter','latex')
ylabel('Residuals (m)','Interpreter','latex')
text(10,23,'(b) between GPS data and optic flow-based control model','Interpreter','latex')
%text(10,15,'(b)','Interpreter','latex')
text(45,13,'0.497 m $\pm$ 2.298 m (mean $\pm$ sd)','Interpreter','latex')
axis([-1 401 -20 20])
box on

print -f6 -dpng -r600 Residuals_magnitude
```
