## Supplementary material for "Optic flow helps explain gulls’ altitude control over seas": data set of gulls takeoffs - Flight modelling: Output_graph_altitude_control_vs_OF_control_ID436.m

```
clc
close all
clear all

load('FlightID436.mat')

altitude1 = altitude(1:length(est_altitude));
clear altitude
altitude = altitude1;
clear altitude1

time1 = time(1:length(est_altitude));
clear time
time = time1;
clear time1

speed1 = speed(1:length(est_altitude));
clear speed
speed = speed1;
clear speed1

cost_func = 'NRMSE';
fit = round(goodnessOfFit(est_altitude,altitude,cost_func)*100);
fitfactor = sprintf('Fit factor: %d%%', fit);
omega = 1/K*180/pi;
%omega_sp = sprintf('Optic flow set-point: %d °/s',omega);
time_constant = round(Tau);
t_c = sprintf('Time constant: %d s',time_constant);
Max_climb = sprintf('V_{Zmax} =  %.2f m/s',Vzmax);
figure(1)
%subplot(3,1,1)
%plot(time,ax,'.b','markersize',15)
%xlabel('Time (s)')
%ylabel('Forward acceleration (m.s^{-2})')

subplot(2,1,1) % figure 2 lignes ; 1 colonne ; 1 N° du graphe
plot(time,speed,'.k','markersize',15)
xlabel('Time (s)')
ylabel('Groundspeed (m.s^{-1})')
axis([0 max(time) 0 25])
title(t,'FontSize',12,'FontWeight','bold')
Tgps = sprintf('Tgps = %.1f s', round(10*Te)/10);
text(0,26,'(a)','FontSize',10,'FontWeight','bold','color','k')
%text(20,20,Tgps,'FontSize',10,'FontWeight','bold','color','k')
text(20,23,'$T_{gps}$ = 14.2 $s$','Interpreter','latex','FontSize',10,'FontWeight','bold','color','k')
grid off

subplot(2,1,2)
plot(time,altitude,'.k','markersize',15)
hold on
plot(time,est_altitude,'ob','markersize',4)
xlabel('Time (s)')
ylabel('Altitude (m)')
grid off
axis([0 max(time) -5 100])
text(0,105,'(b)','FontSize',10,'FontWeight','bold','color','k')
text(300,90,'Optic flow-based control','Interpreter','latex','FontSize',10,'FontWeight','bold','color','b')
text(20,90,'Altitude control','Interpreter','latex','FontSize',10,'FontWeight','bold','color','r')
%text(20,80,fitfactor,'FontSize',10,'FontWeight','bold','color','b')
text(300,80,'Fit factor = 75\%','Interpreter','latex','FontSize',10,'FontWeight','bold','color','b')
text(20,80,'Fit factor = 39\%','Interpreter','latex','FontSize',10,'FontWeight','bold','color','r')
%text(20,70,omega_sp,'FontSize',10,'FontWeight','bold','color','b')
text(300,70,'$\omega_{sp}$ = 17.4 $^\circ/s$','Interpreter','latex','FontSize',10,'FontWeight','bold','color','b')
text(20,70,'$h_{sp}$ = 55.2 $m$','Interpreter','latex','FontSize',10,'FontWeight','bold','color','r')
%text(20,60,t_c,'FontSize',10,'FontWeight','bold','color','b')
text(600,90,'$\tau_{h}$ = 138 $s$','Interpreter','latex','FontSize',10,'FontWeight','bold','color','k')
%text(20,50,Max_climb,'FontSize',10,'FontWeight','bold','color','b')
text(600,80,'$V_{zmax}$ = 0.55 $m/s$','Interpreter','latex','FontSize',10,'FontWeight','bold','color','k')
%fig = sprintf('FlightID%d', ID_flight);
%save(fig,'speed', 'altitude', 'est_altitude','time')

load('FlightID436_alt.mat')
hold on
plot(time,est_altitude,'or','markersize',4)

print('Fig_OF_vs_altitude_ID436','-f1','-dpng','-r600')


return

%% Short movie as illustration
%%

warning off
fId = figure;
writerObj = VideoWriter('fig.avi'); % Name it.
writerObj.FrameRate = 60; % How many frames per second.
open(writerObj); 
gull = imread('gull.png');
figure(fId)
subplot(2,1,1) % figure 2 lignes ; 1 colonne ; 1 N° du graphe
plot(time,speed,'.k','markersize',15)
xlabel('Time (s)')
ylabel('Groundspeed (m.s^{-1})')
axis([0 max(time) 0 25])
title(t,'FontSize',12,'FontWeight','bold')
grid off

subplot(2,1,2)
plot(time,altitude,'.k','markersize',15)
hold on
xlabel('Time (s)')
ylabel('Altitude (m)')
grid off
axis([0 max(time) 0 100])
text(50,90,fitfactor,'FontSize',10,'FontWeight','bold','color','b')
hold on

for i=1:size(est_altitude)      
    % We just use pause but pretend you have some really complicated thing here...
    pause(1);
    figure(fId); % Makes sure you use your desired frame.
    plot(time(i),est_altitude(i),'ob','markersize',4);
    %image(time(i),est_altitude(i),gull) ;
    %if mod(i,4)==0, % Uncomment to take 1 out of every 4 frames.
        frame = getframe(gcf); % 'gcf' can handle if you zoom in to take a movie.
        writeVideo(writerObj, frame);
    %end
 
end
hold off
close(writerObj); % Saves the movie.
```
