## Supplementary material for "Optic flow helps explain gulls’ altitude control over seas": data set of gulls takeoffs - Flight modelling: shadedErrorBar.m

```
function varargout=shadedErrorBar(x,y,errBar,lineProps,transparent)
% function H=shadedErrorBar(x,y,errBar,lineProps,transparent)
%
% Purpose 
% Makes a 2-d line plot with a pretty shaded error bar made
% using patch. Error bar color is chosen automatically.
%
% Inputs
% x - vector of x values [optional, can be left empty]
% y - vector of y values or a matrix of n observations by m cases
%     where m has length(x);
% errBar - if a vector we draw symmetric errorbars. If it has a size
%          of [2,length(x)] then we draw asymmetric error bars with
%          row 1 being the upper bar and row 2 being the lower bar
%          (with respect to y). ** alternatively ** errBar can be a
%          cellArray of two function handles. The first defines which
%          statistic the line should be and the second defines the
%          error bar.
% lineProps - [optional,'-k' by default] defines the properties of
%             the data line. e.g.:    
%             'or-', or {'-or','markerfacecolor',[1,0.2,0.2]}
% transparent - [optional, 0 by default] if ==1 the shaded error
%               bar is made transparent, which forces the renderer
%               to be openGl. However, if this is saved as .eps the
%               resulting file will contain a raster not a vector
%               image. 
%
% Outputs
% H - a structure of handles to the generated plot objects.     
%
%
% Examples
% y=randn(30,80); x=1:size(y,2);
% shadedErrorBar(x,mean(y,1),std(y),'g');
% shadedErrorBar(x,y,{@median,@std},{'r-o','markerfacecolor','r'});    
% shadedErrorBar([],y,{@median,@std},{'r-o','markerfacecolor','r'});    
%
% Overlay two transparent lines
% y=randn(30,80)*10; x=(1:size(y,2))-40;
% shadedErrorBar(x,y,{@mean,@std},'-r',1); 
% hold on
% y=ones(30,1)*x; y=y+0.06*y.^2+randn(size(y))*10;
% shadedErrorBar(x,y,{@mean,@std},'-b',1); 
% hold off
%
%
% Rob Campbell - November 2009
    
%%%%%%%%%%%%%%%%%%%%%%%%%%%%%%%%%%%%%%%%%%%%%%%%%%%%%%%%%%%%%    
% Error checking    
narginchk(3,5)


%Process y using function handles if needed to make the error bar
%dynamically
if iscell(errBar) 
    fun1=errBar{1};
    fun2=errBar{2};
    errBar=fun2(y);
    y=fun1(y);
else
    y=y(:)';
end

if isempty(x)
    x=1:length(y);
else
    x=x(:)';
end


%Make upper and lower error bars if only one was specified
if length(errBar)==length(errBar(:))
    errBar=repmat(errBar(:)',2,1);
else
    s=size(errBar);
    f=find(s==2);
    if isempty(f), error('errBar has the wrong size'), end
    if f==2, errBar=errBar'; end
end

if length(x) ~= length(errBar)
    error('length(x) must equal length(errBar)')
end

%Set default options
defaultProps={'-k'};
if nargin<4, lineProps=defaultProps; end
if isempty(lineProps), lineProps=defaultProps; end
if ~iscell(lineProps), lineProps={lineProps}; end

if nargin<5, transparent=0; end


%%%%%%%%%%%%%%%%%%%%%%%%%%%%%%%%%%%%%%%%%%%%%%%%%%%%%%%%%%%%%    
% Plot to get the parameters of the line 
H.mainLine=plot(x,y,lineProps{:});


% Work out the color of the shaded region and associated lines
% Using alpha requires the render to be openGL and so you can't
% save a vector image. On the other hand, you need alpha if you're
% overlaying lines. There we have the option of choosing alpha or a
% de-saturated solid colour for the patch surface .

col=get(H.mainLine,'color');
edgeColor=col+(1-col)*0.55;
patchSaturation=0.15; %How de-saturated or transparent to make patch
if transparent
    faceAlpha=patchSaturation;
    patchColor=col;
    set(gcf,'renderer','openGL')
else
    faceAlpha=1;
    patchColor=col+(1-col)*(1-patchSaturation);
    set(gcf,'renderer','painters')
end

    
%Calculate the error bars
uE=y+errBar(1,:);
lE=y-errBar(2,:);


%Add the patch error bar
holdStatus=ishold;
if ~holdStatus, hold on,  end


%Make the patch
yP=[lE,fliplr(uE)];
xP=[x,fliplr(x)];

%remove nans otherwise patch won't work
xP(isnan(yP))=[];
yP(isnan(yP))=[];


H.patch=patch(xP,yP,1,'facecolor',patchColor,...
              'edgecolor','none',...
              'facealpha',faceAlpha);


%Make pretty edges around the patch. 
H.edge(1)=plot(x,lE,'-','color',edgeColor);
H.edge(2)=plot(x,uE,'-','color',edgeColor);

%Now replace the line (this avoids having to bugger about with z coordinates)
uistack(H.mainLine,'top')


if ~holdStatus, hold off, end


if nargout==1
    varargout{1}=H;
end
```
