## Supplemental Information for "Optic flow helps explain gulls’ altitude control over seas"

### SI Appendix

| | | | | |
|---|---|---|---|---|
| $h$ | $0$ | | $h_{ref} \cdot \left(\frac{\omega_{sp}}{V_{ref}\cdot\alpha}\right)^{\frac{1}{\alpha-1}}$ | $+\infty$ |
| $\frac{\partial f}{\partial h}$ | | $+$ | $0$ | $-$ |
| $f(h)$ | $V_{air} > 0$ | ↗ | $\hat{f}$ | ↘ $0$ ↘ $-\infty$ |

Table S1: Variation table of the function $f$ described by equation (2.9). There exists only a unique altitude $h$ enabling $f(h) = 0$.

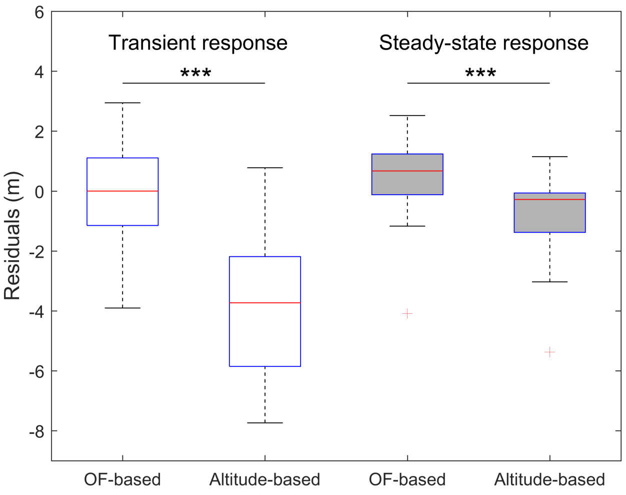


Figure S2: The residuals distribution (in meters) of each model (set of medians in Figs. 8c-d, $n = 27$ for each boxplot) in transient response (white shaded boxes) and in steady-state response (gray shaded boxes). The distributions are presented using classical boxplots showing the median (red line), the 25th and the 75th percentiles (blue box), and the minimum and maximum values (black whiskers). "OF-based" stands for optic flow-based control model and "Altitude-based" stands for altitude control model. The mean of the distribution of the difference between residuals' distributions is significantly higher than the mean altitude error (2.77 m) of the GPS tracker (Bouten *et al.* 2013) in transient response (one-sided $t$-test, $n = 27$, $p \ll 0.001$).

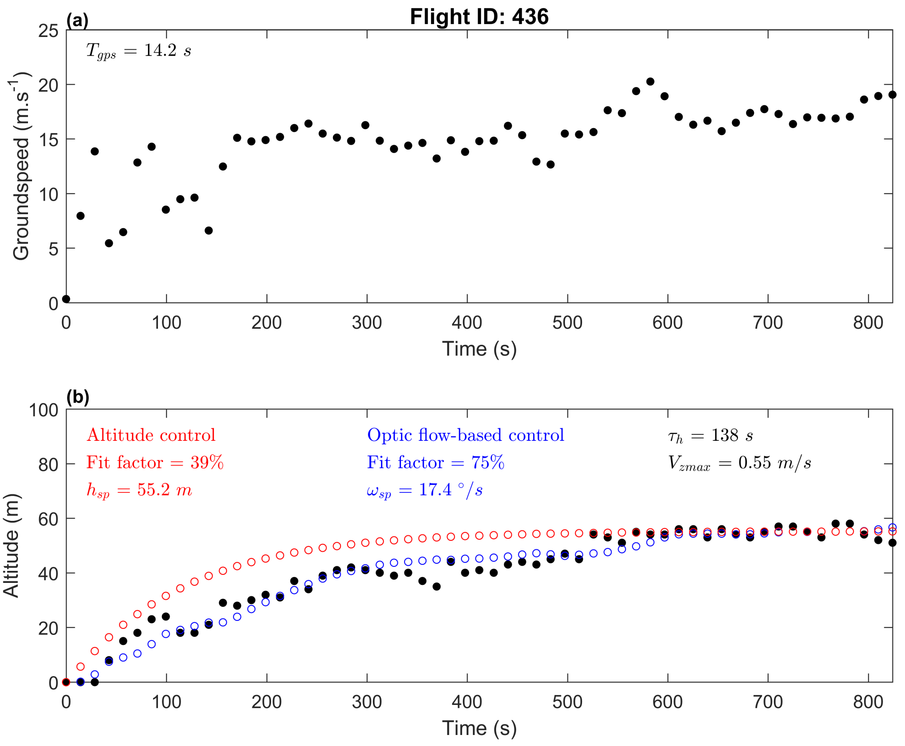


Figure S3: (a) Groundspeed of the gull ID426 tracked with the GPS. (b) Altitude of the gull: black dots represent the GPS data, red dots represent the gull altitude on the basis of an altitude-based control model (fit factor: 39%), and the blue dots represent the gull altitude on the basis of an optic flow-based control model (fit factor: 75%). A significant correlation was observed between groundspeed and altitude of the GPS data ($\rho = 0.71$, $p \ll 0.001$ by Spearman's test).

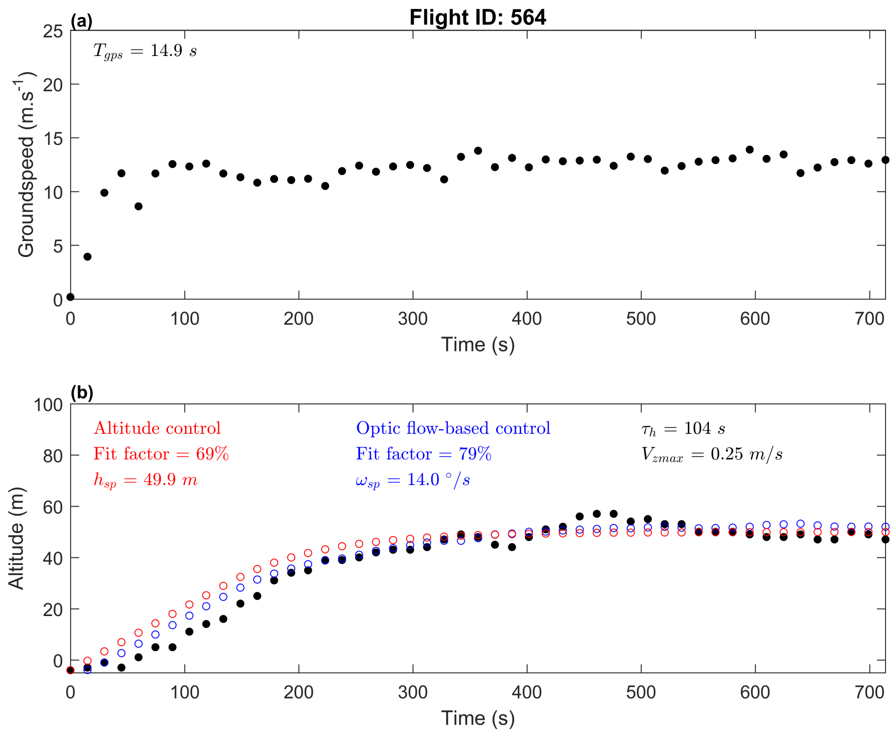


Figure S4: (a) Groundspeed of the gull ID564 tracked with the GPS. (b) Altitude of the gull: black dots represent the GPS data, red dots represent the gull altitude on the basis of an altitude-based control model (fit factor: 69%), and the blue dots represent the gull altitude on the basis of an optic flow-based control model (fit factor: 79%). A significant correlation was observed between groundspeed and altitude of the GPS data ($\rho = 0.68$, $p \ll 0.001$ by Spearman's test).

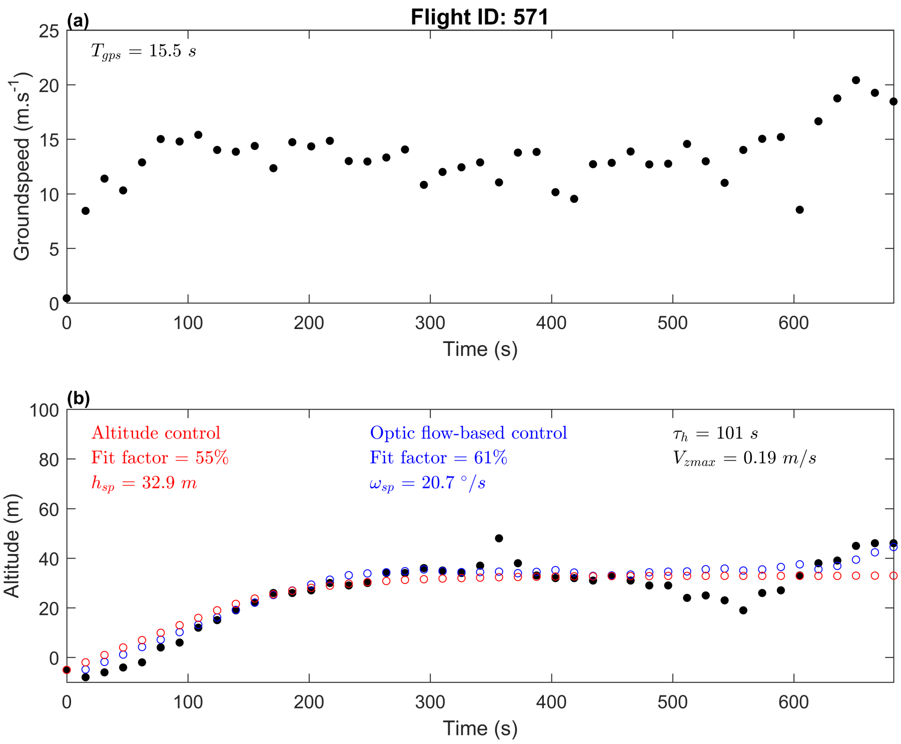


Figure S6:(a) Groundspeed of the gull ID571 tracked with the GPS. (b) Altitude of the gull: black dots represent the GPS data, red dots represent the gull altitude on the basis of an altitude-based control model (fit factor: 55%), and the blue dots represent the gull altitude on the basis of an optic flow-based control model (fit factor: 61%). No correlation was observed between groundspeed and altitude of the GPS data.

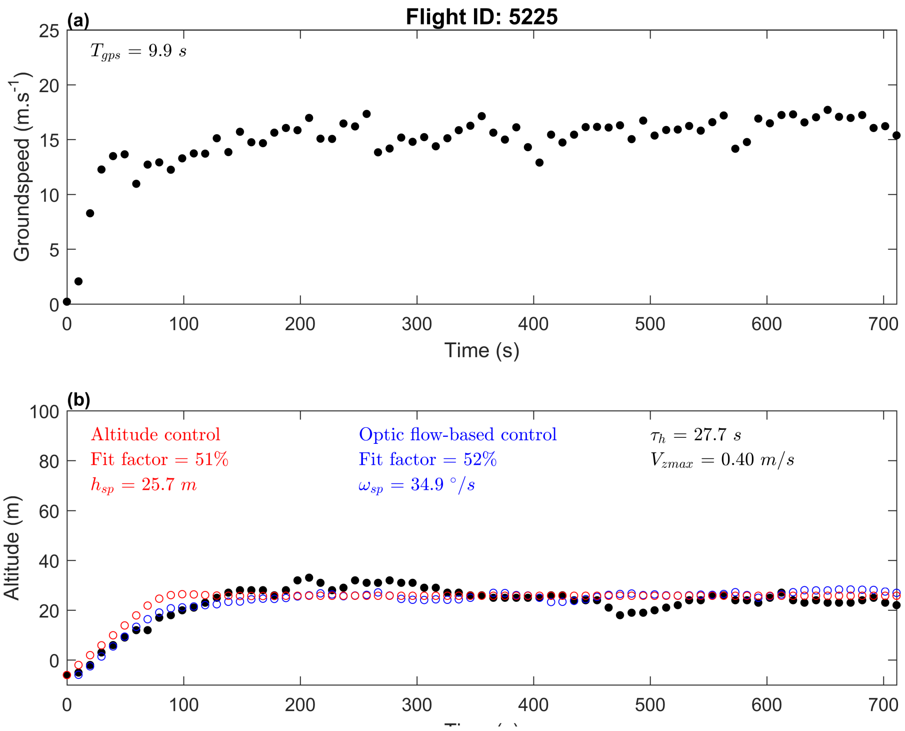


Figure S6:(a) Groundspeed of the gull ID5225 tracked with the GPS. (b) Altitude of the gull: black dots represent the GPS data, red dots represent the gull altitude on the basis of an altitude-based control model (fit factor: 51%), and the blue dots represent the gull altitude on the basis of an optic flow-based control model (fit factor: 52%). A significant correlation was observed between groundspeed and altitude of the GPS data ($\rho = 0.27$, $p \ll 0.001$ by Spearman's test).

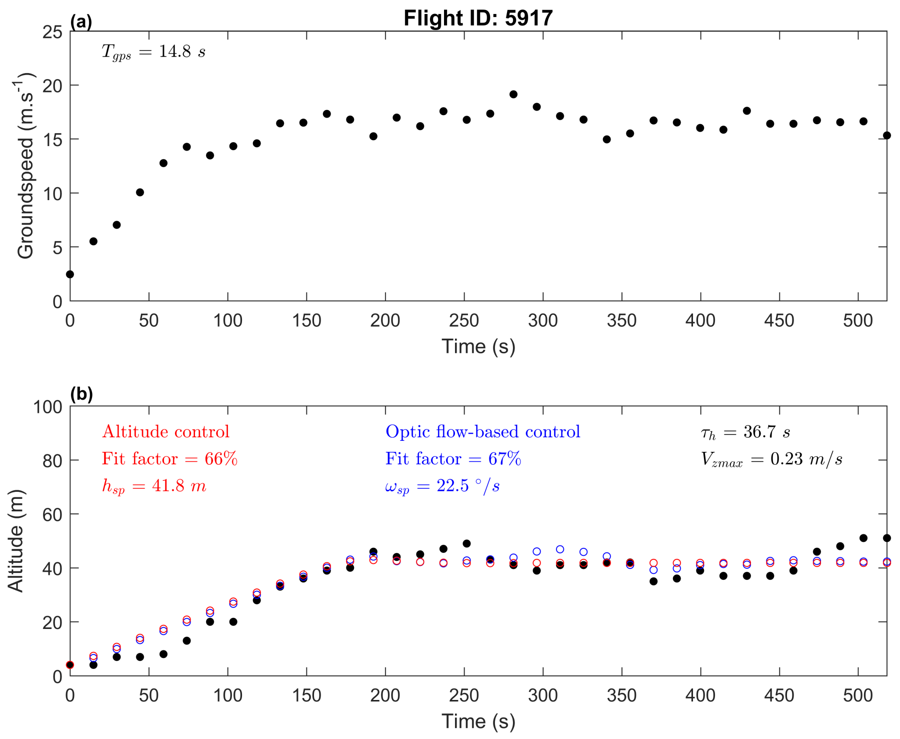


Figure S7:(a) Groundspeed of the gull ID5917 tracked with the GPS. (b) Altitude of the gull: black dots represent the GPS data, red dots represent the gull altitude on the basis of an altitude-based control model (fit factor: 66%), and the blue dots represent the gull altitude on the basis of an optic flow-based control model (fit factor: 67%). A significant correlation was observed between groundspeed and altitude of the GPS data ($\rho = 0.57$, $p \ll 0.001$ by Spearman's test).

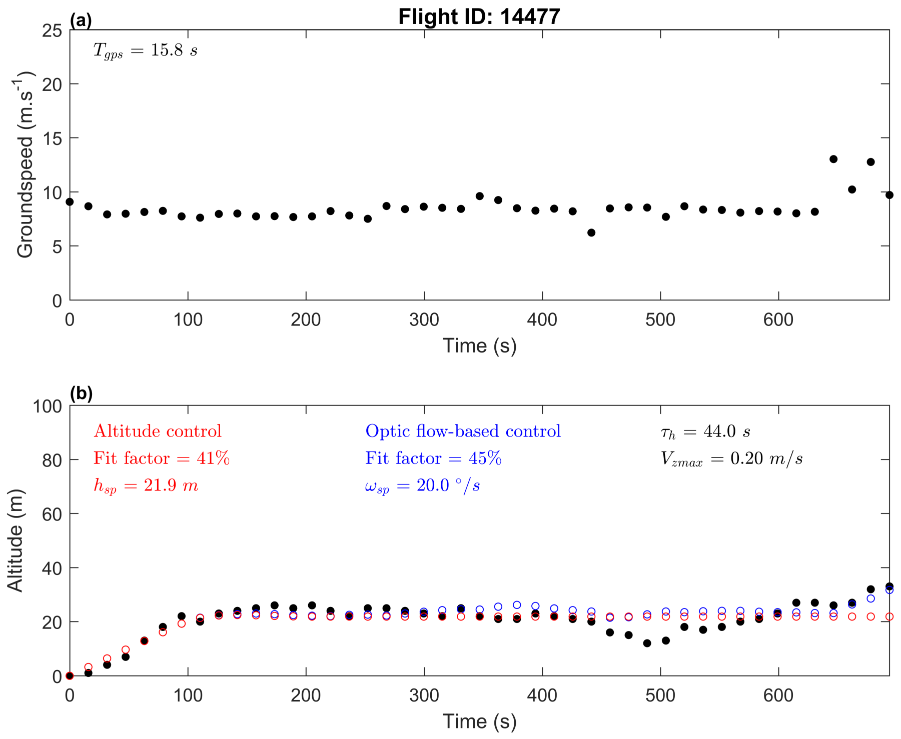


Figure S8:(a) Groundspeed of the gull ID14477 tracked with the GPS. (b) Altitude of the gull: black dots represent the GPS data, red dots represent the gull altitude on the basis of an altitude-based control model (fit factor: 41%), and the blue dots represent the gull altitude on the basis of an optic flow-based control model (fit factor: 45%). No significant correlation was observed between groundspeed and altitude of the GPS data.

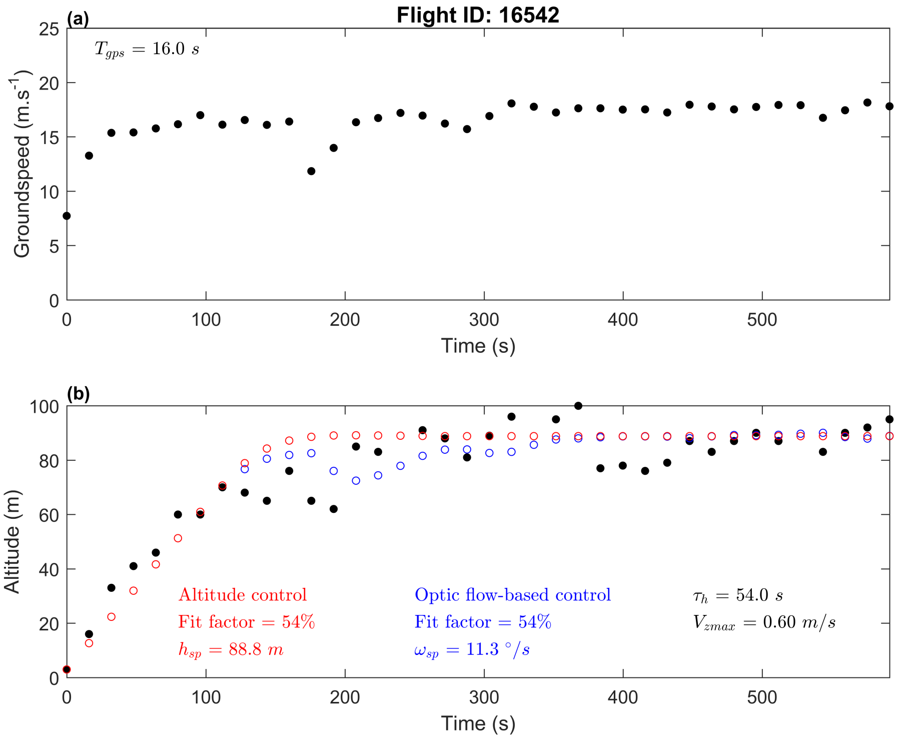


Figure S19:(a) Groundspeed of the gull ID16542 tracked with the GPS. (b) Altitude of the gull: black dots represent the GPS data, red dots represent the gull altitude on the basis of an altitude-based control model (fit factor: 54%), and the blue dots represent the gull altitude on the basis of an optic flow-based control model (fit factor: 54%). No significant correlation was observed between groundspeed and altitude of the GPS data.

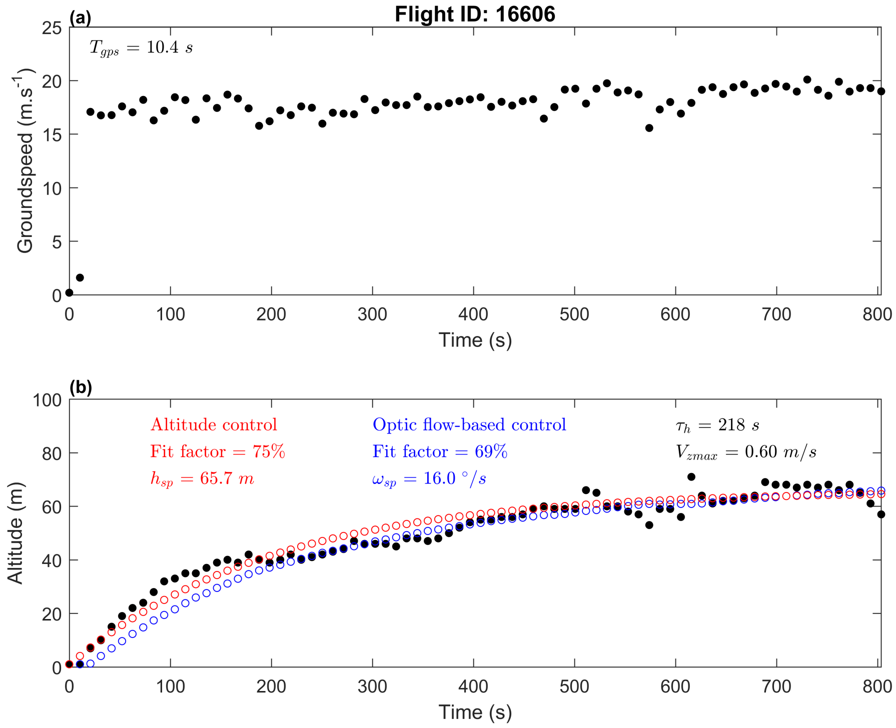


Figure S10:(a) Groundspeed of the gull ID16606 tracked with the GPS. (b) Altitude of the gull: black dots represent the GPS data, red dots represent the gull altitude on the basis of an altitude-based control model (fit factor: 75%), and the blue dots represent the gull altitude on the basis of an optic flow-based control model (fit factor: 69%). A significant correlation was observed between groundspeed and altitude of the GPS data ($\rho = 0.71$, $p \ll 0.001$ by Spearman's test).

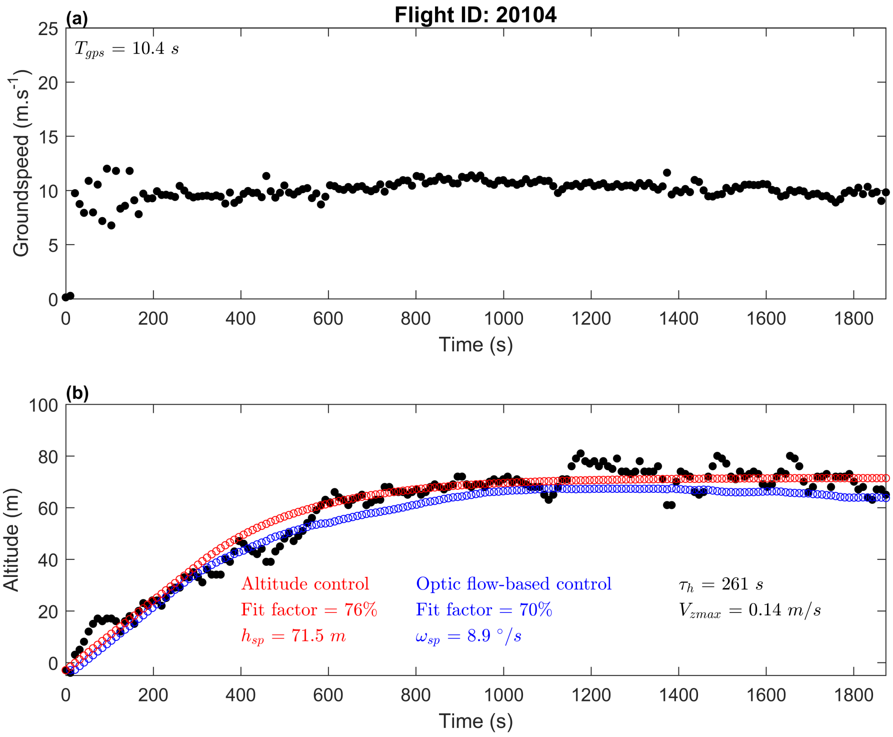


Figure S11:(a) Groundspeed of the gull ID20104 tracked with the GPS. (b) Altitude of the gull: black dots represent the GPS data, red dots represent the gull altitude on the basis of an altitude-based control model (fit factor: 76%), and the blue dots represent the gull altitude on the basis of an optic flow-based control model (fit factor: 70%). A significant correlation was observed between groundspeed and altitude of the GPS data ($\rho = 0.35$, $p \ll 0.001$ by Spearman's test).

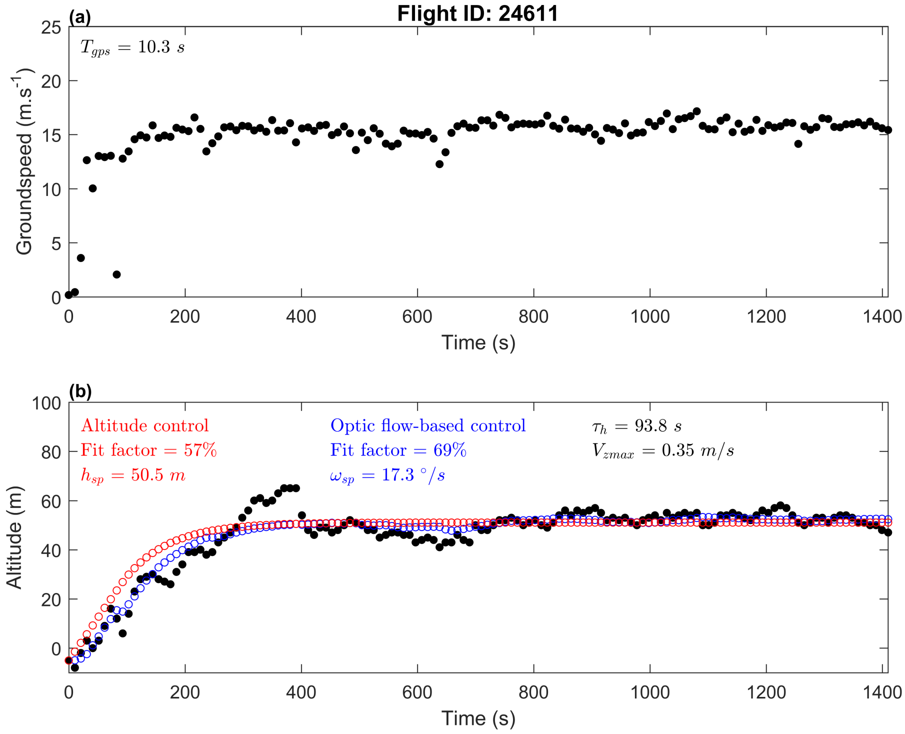


Figure S12:(a) Groundspeed of the gull ID24611 tracked with the GPS. (b) Altitude of the gull: black dots represent the GPS data, red dots represent the gull altitude on the basis of an altitude-based control model (fit factor: 57%), and the blue dots represent the gull altitude on the basis of an optic flow-based control model (fit factor: 69%). A significant correlation was observed between groundspeed and altitude of the GPS data ($\rho = 0.51$, $p \ll 0.001$ by Spearman's test).

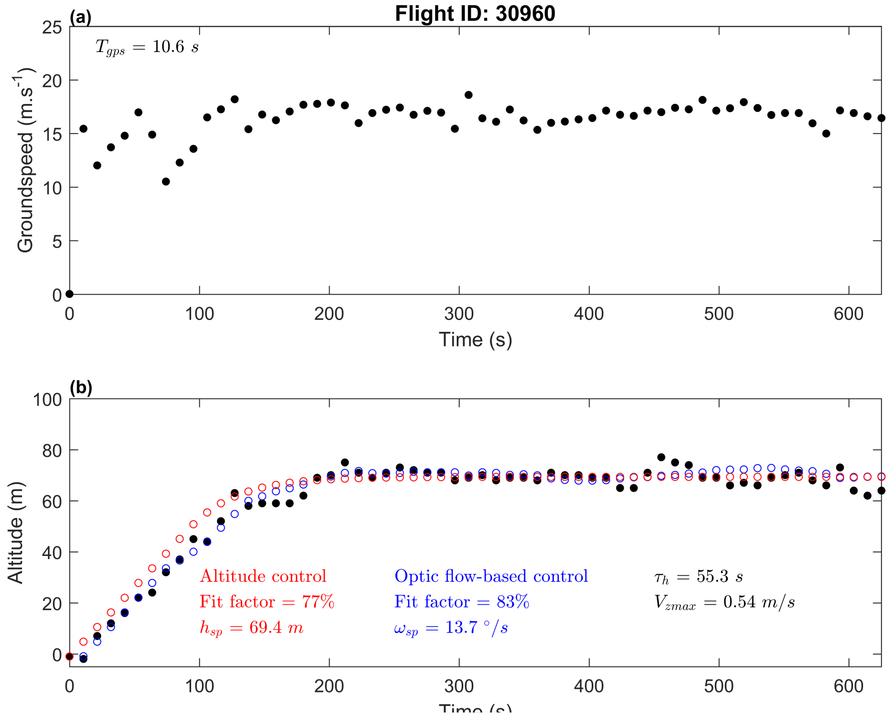


Figure S13:(a) Groundspeed of the gull ID30960 tracked with the GPS. (b) Altitude of the gull: black dots represent the GPS data, red dots represent the gull altitude on the basis of an altitude-based control model (fit factor: 77%), and the blue dots represent the gull altitude on the basis of an optic flow-based control model (fit factor: 83%). A significant correlation was observed between groundspeed and altitude of the GPS data ($\rho = 0.48$, $p \ll 0.001$ by Spearman's test).

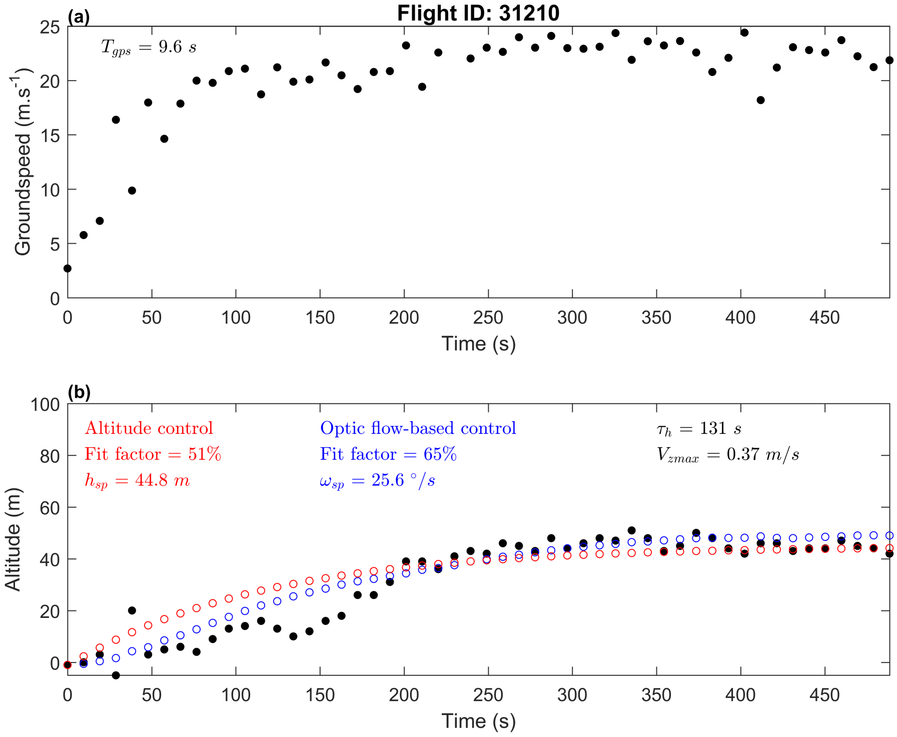


Figure S14:(a) Groundspeed of the gull ID31210 tracked with the GPS. (b) Altitude of the gull: black dots represent the GPS data, red dots represent the gull altitude on the basis of an altitude-based control model (fit factor: 51%), and the blue dots represent the gull altitude on the basis of an optic flow-based control model (fit factor: 65%). A significant correlation was observed between groundspeed and altitude of the GPS data ($\rho = 0.70$, $p \ll 0.001$ by Spearman's test).

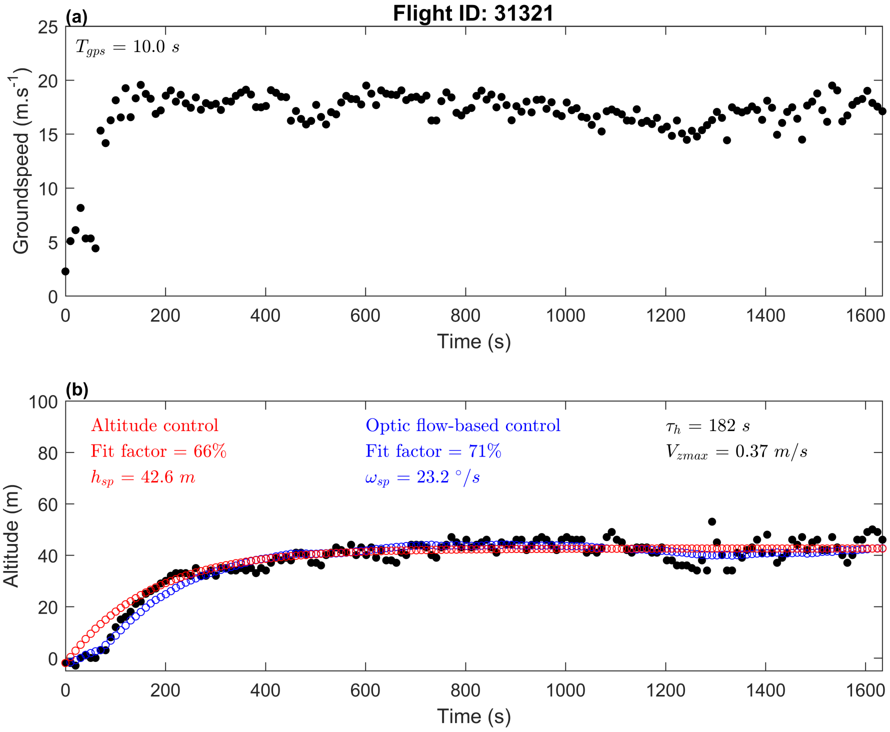


Figure S15:(a) Groundspeed of the gull ID31321 tracked with the GPS. (b) Altitude of the gull: black dots represent the GPS data, red dots represent the gull altitude on the basis of an altitude-based control model (fit factor: 66%), and the blue dots represent the gull altitude on the basis of an optic flow-based control model (fit factor: 71%). A significant correlation was observed between groundspeed and altitude of the GPS data ($\rho = 0.22$, $p \ll 0.001$ by Spearman's test).

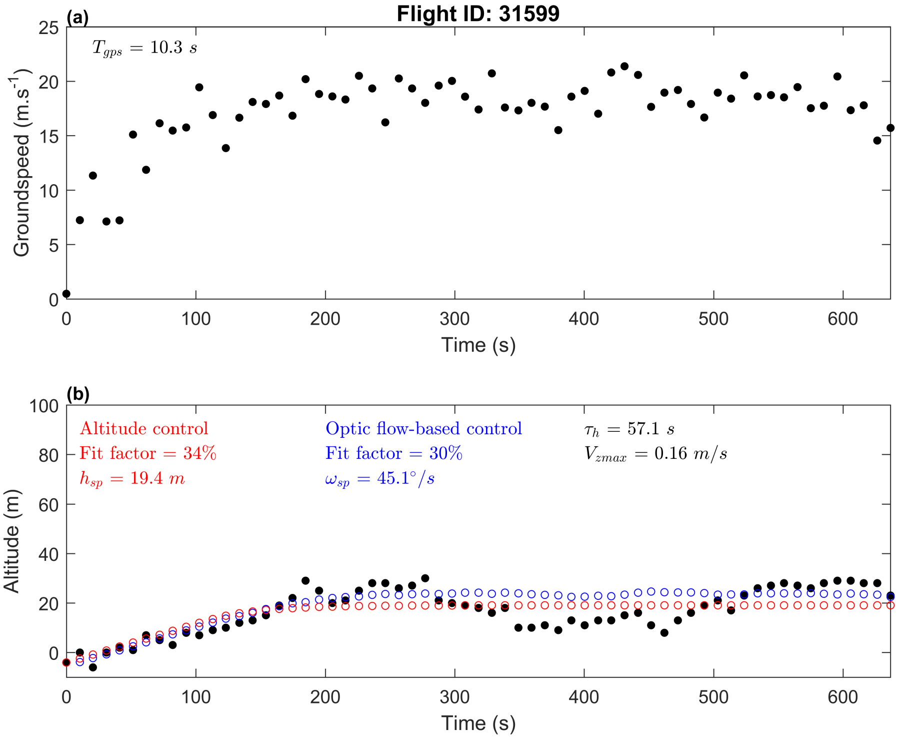


Figure S16:(a) Groundspeed of the gull ID31599 tracked with the GPS. (b) Altitude of the gull: black dots represent the GPS data, red dots represent the gull altitude on the basis of an altitude-based control model (fit factor: 34%), and the blue dots represent the gull altitude on the basis of an optic flow-based control model (fit factor: 30%). A significant correlation was observed between groundspeed and altitude of the GPS data ($\rho = 0.46$, $p \ll 0.001$ by Spearman's test).

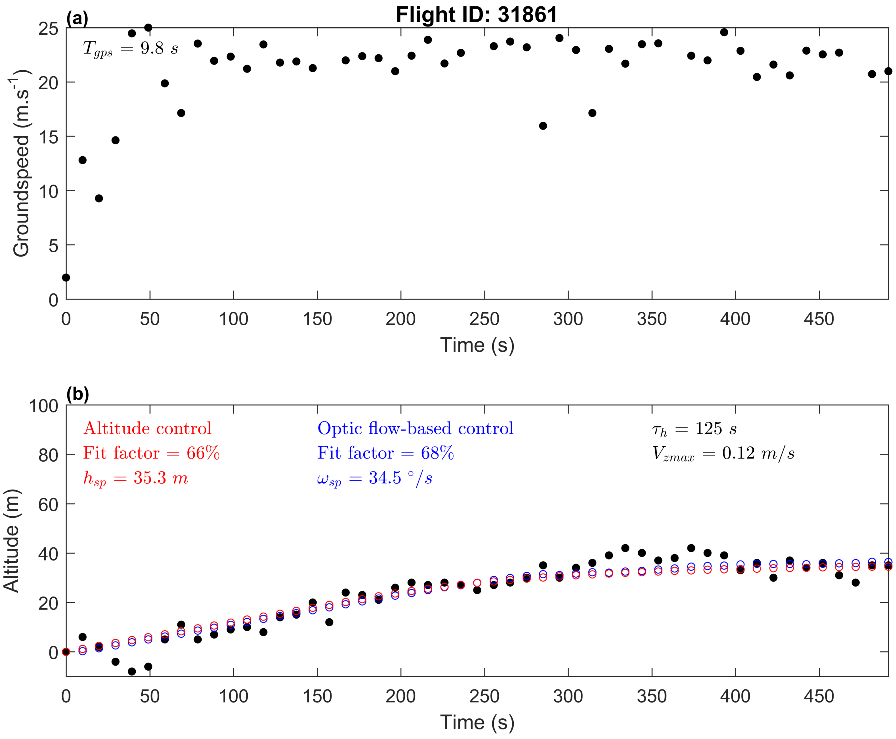


Figure S17:(a) Groundspeed of the gull ID31861 tracked with the GPS. (b) Altitude of the gull: black dots represent the GPS data, red dots represent the gull altitude on the basis of an altitude-based control model (fit factor: 66%), and the blue dots represent the gull altitude on the basis of an optic flow-based control model (fit factor: 68%). No significant correlation was observed between groundspeed and altitude of the GPS data.

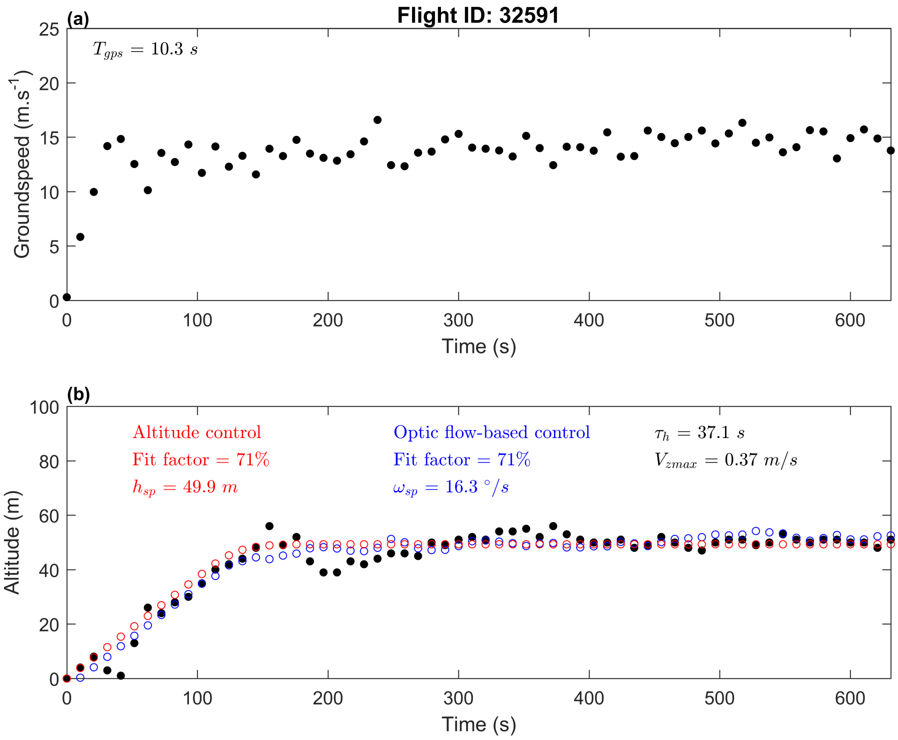


Figure S18:(a) Groundspeed of the gull ID32591 tracked with the GPS. (b) Altitude of the gull: black dots represent the GPS data, red dots represent the gull altitude on the basis of an altitude-based control model (fit factor: 71%), and the blue dots represent the gull altitude on the basis of an optic flow-based control model (fit factor: 71%). No significant correlation was observed between groundspeed and altitude of the GPS data.

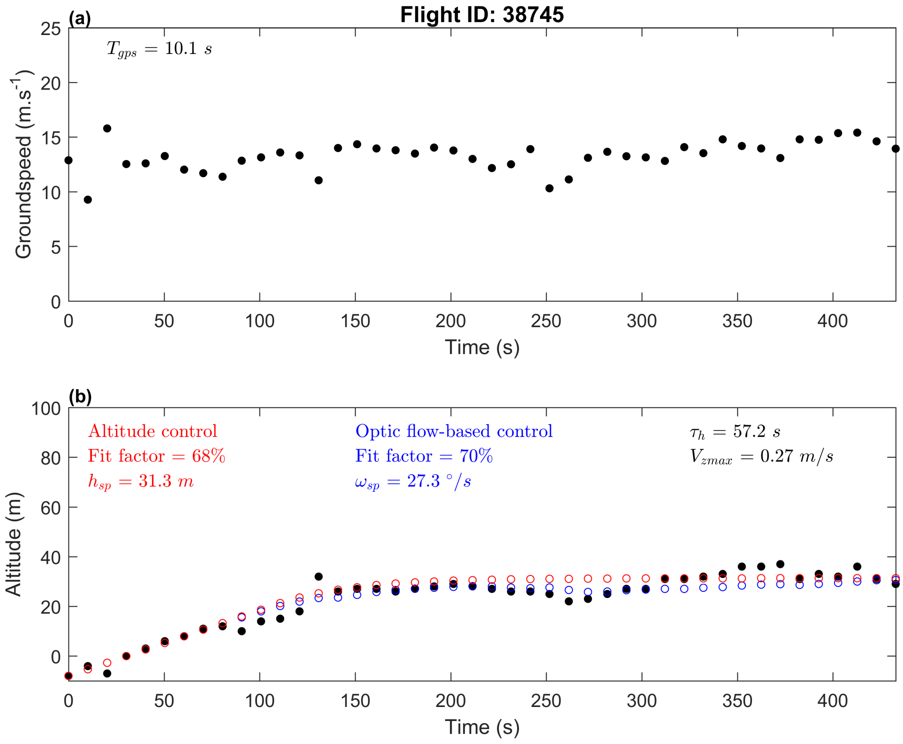


Figure S19:(a) Groundspeed of the gull ID38745 tracked with the GPS. (b) Altitude of the gull: black dots represent the GPS data, red dots represent the gull altitude on the basis of an altitude-based control model (fit factor: 68%), and the blue dots represent the gull altitude on the basis of an optic flow-based control model (fit factor: 70%). A significant correlation was observed between groundspeed and altitude of the GPS data ($\rho = 0.51$, $p \ll 0.001$ by Spearman's test).

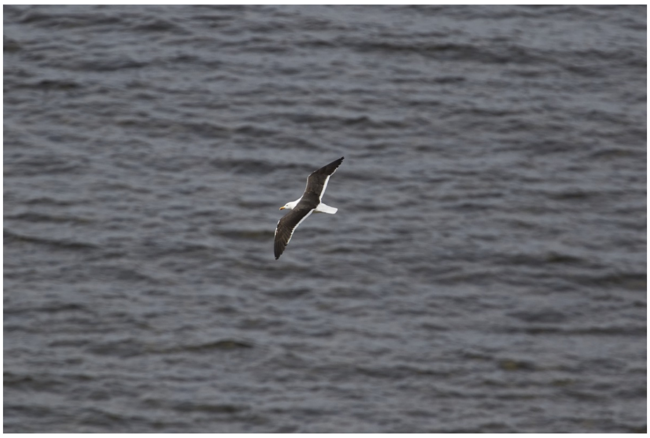

Figure S20: Gull (*Larus fuscus*) flying above the Baltic sea. The state of the sea while gulls are flying is typically with a Beaufort number of 3 to 4. Photographic credit and courtesy from Aron Hejdström.
